# Amphetamine disrupts dopamine axon growth in adolescence by a sex-specific mechanism

**DOI:** 10.1101/2022.12.14.520468

**Authors:** Lauren M. Reynolds, Giovanni Hernandez, Christina Popescu, Del MacGowan, Dominique Nouel, Santiago Cuesta, Samuel Burke, Katherine E. Savell, Janet Zhao, Jose Maria Restrepo-Lozano, Matthew Pokinko, Michel Giroux, Sonia Israel, Taylor Orsini, Susan He, Michael Wodzinski, Julia G. Epelbaum, Louis-Éric Trudeau, Bryan Kolb, Jeremy J. Day, Cecilia Flores

## Abstract

Initiating drug use during adolescence increases the risk of developing addiction and psychiatric disorders later in life, with long-term outcomes varying according to sex and exact timing of use. Even though most individuals begin experimenting with drugs of abuse in adolescence, to date, the cellular and molecular underpinnings explaining differential sensitivity to detrimental drug effects remain unknown. The Netrin-1/DCC guidance cue system plays a critical role in the adolescent development of mesocorticolimbic dopamine circuitry, segregating the cortical and limbic pathways. Adolescent experiences, including exposure to drugs of abuse, can regulate *Dcc* expression in male mice, placing Netrin-1/DCC signaling as a potential molecular link between experience and enduring changes to circuitry and behavior. Here we show that exposure to a recreational-like regimen of amphetamine (AMPH) in adolescence induces sex- and age-specific alterations in *Dcc* expression in the ventral tegmental area. Female mice are protected against the deleterious long-term effects of AMPH-induced *Dcc* regulation by compensatory changes in the expression of its binding partner, Netrin-1. AMPH induces targeting errors in mesolimbic dopamine axons and triggers their ectopic growth to the prefrontal cortex, only in early-adolescent male mice, underlying a male-specific vulnerability to its enduring cognitive effects. Upregulating DCC receptor expression in dopamine neurons in adolescent males using a neuron-optimized CRISPR/dCas9 Activation System induces female-like protection against the persistent effects of AMPH in early adolescence on inhibitory control. Netrin-1/DCC signaling is therefore a molecular switch which can be differentially regulated in response to the same experience as function of age and sex of the individual, leading to divergent long-term outcomes associated with vulnerable or resilient phenotypes.

## Introduction

Adolescence is a conserved period of life, encompassing the gradual transition from a juvenile to an adult state. While best characterized in humans,^1^ significant behavioral and neurobiological changes also demarcate an adolescent period in other mammals, including rodents.^2–5^ The dopamine neurotransmitter system continues to mature into adulthood, in humans,^6–10^ non-human primates,^11,12^ and in rodents;^4,13,14^ undergoing robust changes in connectivity and function during adolescence. Because dopamine circuitry development is highly shaped by ongoing experiences in adolescence, it is increasingly conceptualized as a “plasticity system”.^15^ However, the molecular mechanisms by which experiences in adolescence modify dopamine development and enduringly alter its function remain a topic of intense research.

Adolescent experiences with significant neurodevelopmental consequences range from essential/formative (e.g. social interactions with peers or conspecifics)^16–21^ and enriching (e.g. targeted diet and exercise),^22–29^ to deleterious (e.g. excessive stress, bullying).^30–35^ However, one of the experiences that leaves the most lasting mark on the adolescent brain is exposure to drugs of abuse,^36,37^ which epidemiological evidence indeed associates with a lifelong increase in the risk for addiction.^38–42^ While addiction and substance use disorders were once thought to disproportionately affect men, the prevalence of drug abuse in women and in adolescent girls has dramatically increased,^43,44^ highlighting the urgent need to consider both sexes in clinical and preclinical research projects.^45^ Earlier adolescent age of onset of drug use is a powerful predictor of addiction risk in both sexes,^38,39,41^ but marked sex differences in addiction trajectories also exist, with patterns of transition from recreational to compulsive drug use differing between men and women.^46–49^ It is clear that not all adolescents face the same drug exposure on equal footing, and the mechanisms that explain how age and sex modulate the long-term effects of adolescent drug use need to be elucidated.

Guidance cues, widely studied in the context of embryonic growth,^50–52^ have emerged as key organizers of adolescent dopamine development.^4,13,14^ In particular, the Netrin-1/DCC guidance cue system sculpts the structural and functional organization of the mesocorticolimbic dopamine pathway in adolescence by actively segregating the mesocortical and mesolimbic dopamine pathways at the level of the nucleus accumbens. This region functions as a choice point for dopamine axons to remain there and undergo DCC-dependent targeting processes, or to instead to continue growing to the prefrontal cortex.^53,54^ Even subtle disruption to the establishment of mesocorticolimbic dopamine connectivity in adolescence, which can be induced by modifications in Netrin-1 or DCC expression, produce persistent dysregulation of prefrontal pyramidal neuronal structure and function. These enduring changes to the prefrontal cortex result in lasting impairments in impulse control,^53,55^ notably in action inhibition – a known index of vulnerability for addiction.^56,57^ Evidence suggests that experiences in adolescence regulate Netrin-1 and DCC expression, but whether this regulation produces enduring impulse control deficits as a function of both the timing of the experience and the sex of the animal remains unknown. Here we combined molecular, anatomical, and behavioral analysis with targeted gene activation experiments in mice to show that the differential regulation of the Netrin-1/DCC guidance cue system by the same drug experience in adolescence encodes sex- and age-specific consequences on proximal dopamine development and on long-term cognitive outcomes.

## Results

### Experience with amphetamine in adolescence sex-specifically regulates *Dcc* expression in dopamine neurons

The guidance cue receptor DCC and its microRNA repressor miR-218 are highly enriched in dopamine neurons of the rodent ventral tegmental area (VTA; Fig 1a,b).^58,59^ The expression of DCC protein and *Dcc* mRNA in the VTA decreases across postnatal development,^58,60,61^ and can be altered by experience at discrete time points. Whether the effects of experience on *Dcc* expression are both age- and sex-specific remains to be explored. To address this question, we treated male and female mice with recreational-like doses of amphetamine (AMPH; 4 mg/kg; which produces similar plasma levels in mice as recreational exposure of d-amphetamine in humans, including adolescents) ^62^ or saline during early adolescence and quantified *Dcc* mRNA one week later (Fig 1c). Since in males AMPH in early adolescence downregulated *Dcc* mRNA in the VTA by upregulating miR-218,^58^ we also quantified miR-218 expression. Using sex as a biological variable and treatment as factors, the analysis revealed that AMPH in early adolescence downregulates significantly *Dcc* mRNA in the VTA of males, but not in females (Fig 1d). The upregulation of miR-218 levels by AMPH in early adolescent males, which mediates the effects of AMPH on *Dcc* mRNA expresssion,^58^ is not evident in females (Fig 1e). Furthermore, the negative relationship between *Dcc* and miR-218 levels in males in early adolescence (Fig 1f)^58^ is notably absent in females (Fig 1g).

**Figure 1.**
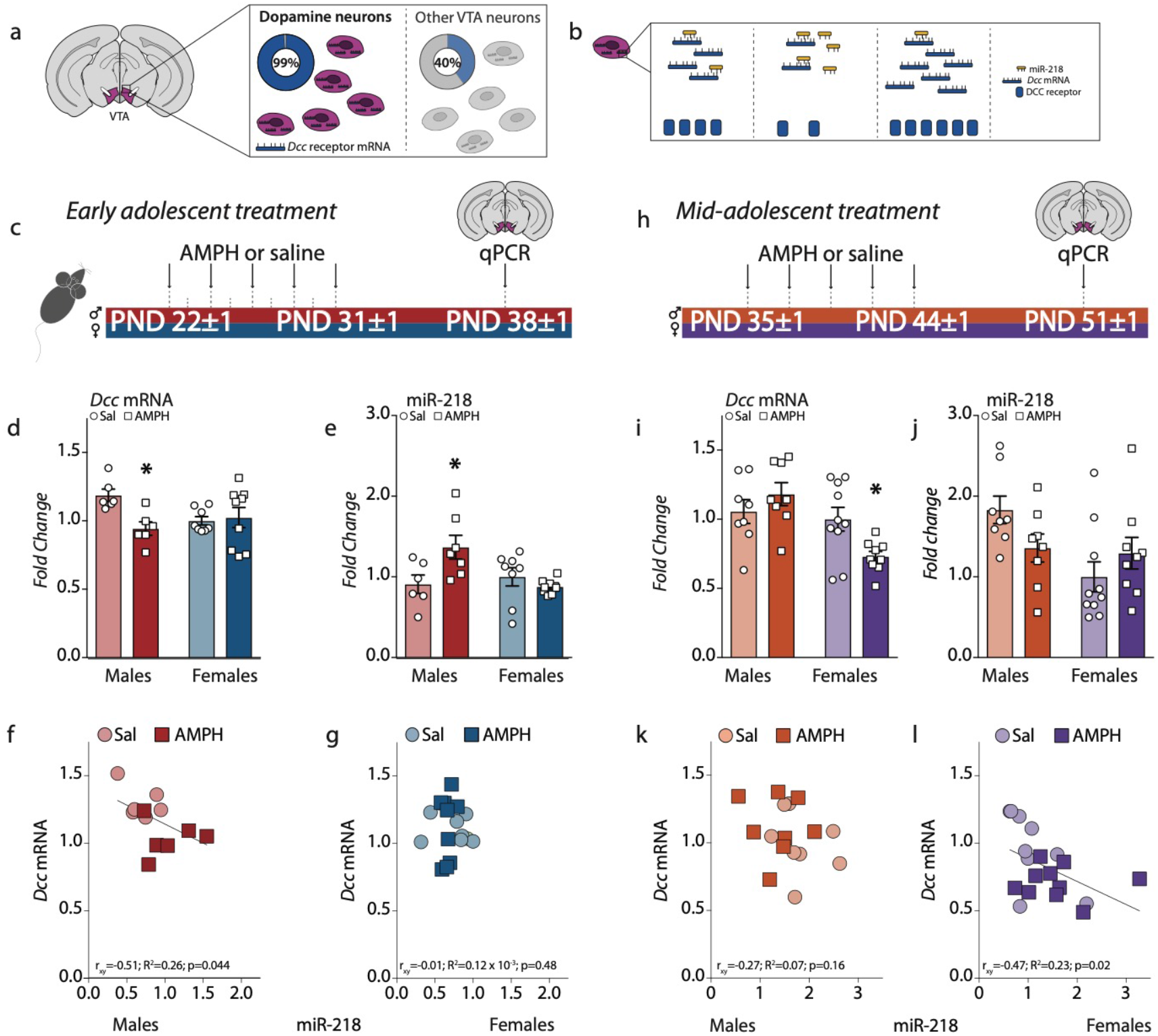
Regulation of Dcc expression in the VTA by AMPH in adolescence is sexually dimorphic. (a) *Dcc* mRNA is expressed by 99% of dopamine neurons in the VTA.^59^ (b) The microRNA miR-218 represses *Dcc* mRNA expression.^58,86^ (c) Timeline of experiments in early adolescence. Male and female mice were exposed to a recreational-like amphetamine (AMPH, 4 mg/kg) regimen from P21±1 to P31±1.^62^ One week later, *Dcc* mRNA and miR-218 expression was quantified in the VTA using qPCR. (d-j) AMPH in early adolescence downregulated *Dcc* expression in males, but not females (d) and increased miR-218 only in males (e) (Table 1A,B). In early adolescence, VTA miR-218 and *Dcc* mRNA levels correlated positively in male, (f) but not female mice (g) (Table 1C,D). (h) Timeline of experiments in mid-adolescence. Male and female mice were exposed to the same recreational-like AMPH regimen, but from P35±1 to P44±1 and VTA transcripts were quantified one week later. (i-l) In mid-adolescence, AMPH no longer altered *Dcc* mRNA in males but downregulated levels in females (i), and it did not significantly alter miR-218 expression in either group (j) (Table 1E,F). In mid-adolescence, VTA miR-218 and *Dcc* mRNA levels did not correlate in males (k) but were positively associated in females (l) (Table 1G,H). Graphs are normalized to the saline condition in female mice (d,e,i,j).

We next exposed male and female mice to the same AMPH treatment regimen, but this time during mid-adolescence (Fig 1h), when we have previously seen an inability of AMPH to regulate DCC receptor expression in males.^63^ We find that AMPH in mid-adolescence also produces a sex-specific effect, but interestingly in the *opposite* direction to what we observed in early adolescence: *Dcc* mRNA in the VTA is downregulated by AMPH in females only (Fig 1i). In addition, there is a treatment by sex interaction in miR-218 expression (Fig 1j), and while no relationship is apparent between *Dcc* mRNA and miR-218 in mid-adolescent males (Fig 1k), these transcripts are significantly and negatively correlated in mid-adolescent females (Fig 1l). These results identify *Dcc* in the VTA as the first developmental genetic marker shown to be regulated by experience in adolescence in a sexually dimorphic manner.

**Table 1:**
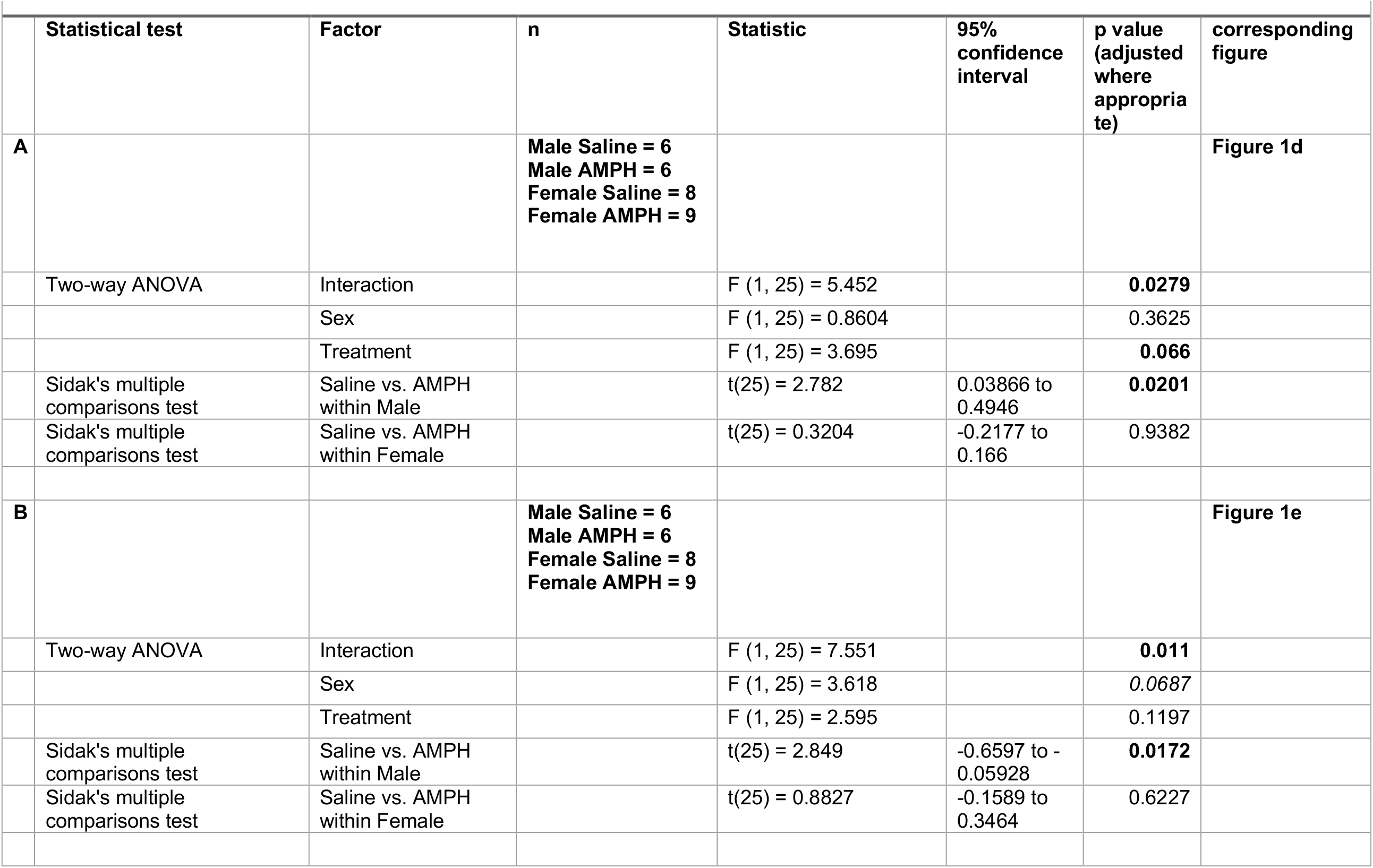
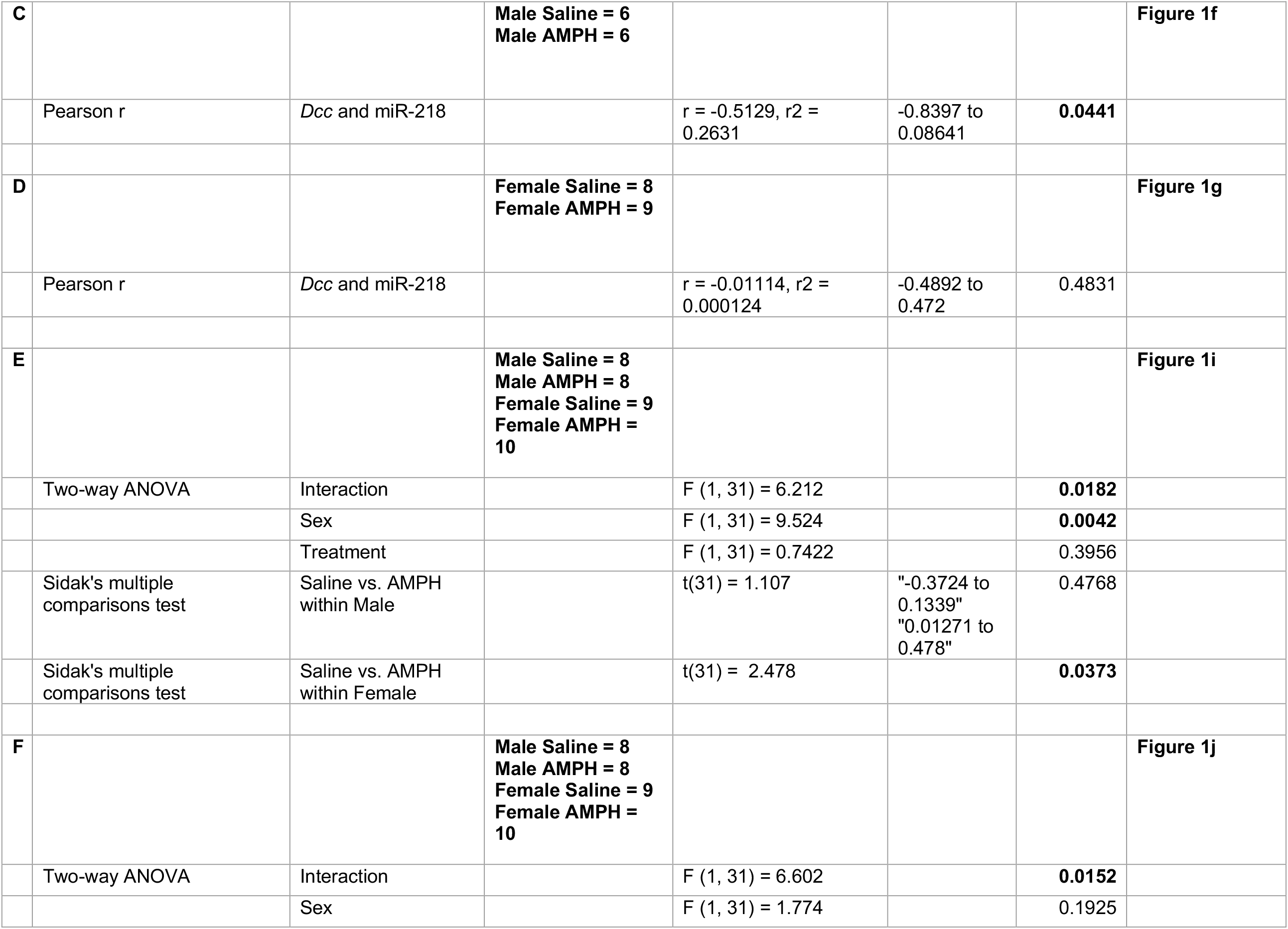
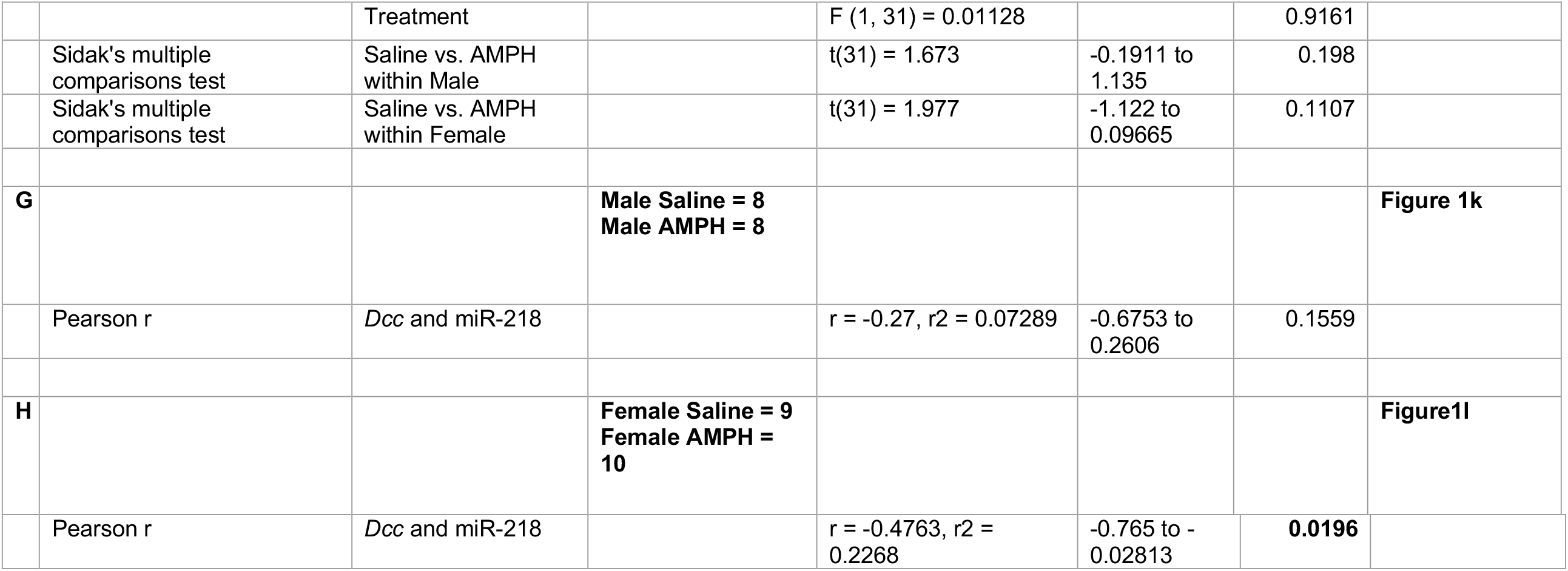
Detailed statistics for Figure 1.

### Female mice are protected from the enduring effects of AMPH in mid-adolescence, despite the downregulation of *Dcc* in dopamine neurons

Our body of work has linked AMPH exposure during early adolescence to enduring changes in prefrontal cortex (PFC) dopamine structure and impulse control in male mice.^54,62,64,65^ These effects are restricted to early adolescence and coincide with the ability of AMPH to downregulate *Dcc* expression.^65^ Since AMPH does not downregulate *Dcc* expression in the VTA of female mice at this adolescent age, we hypothesized that it would not lead to enduring changes in PFC dopamine innervation or impulsivity. However, we predicted that in mid-adolescence, when AMPH does downregulate *Dcc* in females, there would be aberrant dopamine innervation to the PFC and impairments in inhibitory control. Female mice were therefore again exposed to AMPH or saline during early or mid-adolescence (Fig 2a), we found that this regimen indeed produces robust conditioned place preference (Extended data Fig 2a, e). In adulthood, mice were randomly assigned to experiments to either stereologically assess the expanse of the dopamine innervation to the PFC, or to test behavioral inhibition using a Go-No/Go task (Fig 2b). In line with our predictions, and in stark contrast to our findings in males,^64^ AMPH in early adolescent females does not increase the span of dopamine innervation in the PFC (Fig 2c).

**Figure 2.**
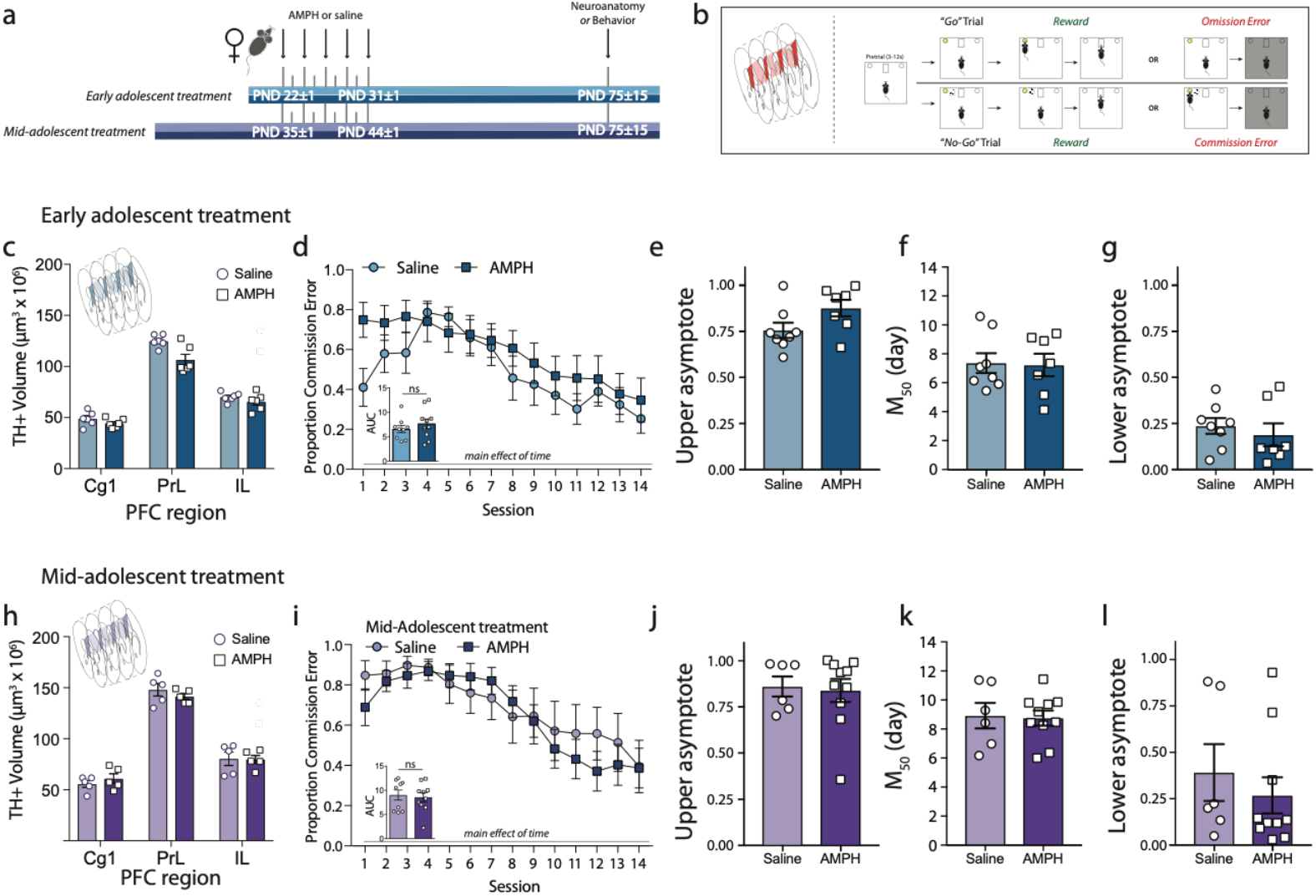
Females are protected against the detrimental effects of AMPH in adolescence on the maturation of PFC dopamine connectivity and impulse control. (a) Timeline for the experiments conducted in female mice treated with the recreational-like amphetamine (AMPH, 4 mg/kg) regimen from P21±1 to P31±1 or from P35±1 to P45±1. (b) In adulthood, mice were randomly assigned to have their brains processed for stereological quantification of PFC dopamine innervation *(left schematic)* or were tested for impulse control in the Go/No-Go task *(right schematic)*. During the Go/No-Go task mice have to respond to a light cue to receive a food reward, but to withhold their response when a tone is paired with the light cue. (c) AMPH in early adolescence does not augment the span of the dopamine input to the cingulate (Cg1), prelimbic (PrL), and infralimbic (IL) subregions of the medial PFC in female mice, in contrast to our previous results in males.^54,64^ Instead, a decrease in the volume of dopamine input to the PrL is evident (Table 2A). (d) AMPH in early adolescence does not alter action impulsivity in adulthood in female mice, unlike our previous observations in males.^65^ Area under the curve (AUC) analysis indicates that the proportion of commission errors is similar between the AMPH-treated and saline groups (Table 2B-C). (e-g) Sigmoidal curve fit analysis (Table 2D-F) further revealed that there is no difference in the number of commission errors at the beginning of the task (e, upper asymptote), that both groups began to improve their inhibitory control performance around day 7-8 (f, M50), and that both groups show similar proportion of commission errors during the last sessions (g, lower asymptote). (h) AMPH in mid-adolescence does not alter the extent of the dopamine input to the Cg1, PrL and IL subregions of the PFC in female mice (Table 2G), despite downregulating *Dcc* in dopamine neurons (Figure 1i). (i-l) In mid-adolescence, females continue to be insensitive to AMPH-induced deficits in action impulsivity, with no differences in the proportion of commission errors in the task (Table 2H-I). (i) Sigmoidal curve fit analysis (Table 2J-L) revealed that AMPH and saline groups perform equally at the beginning of the task (j, upper asymptote), start showing improvement around day 7-8 (j, M50), and have similar low proportion of commission errors during the last session (l, lower asymptote).

Furthermore, while PFC dopamine disruptions produced by AMPH in early adolescence associate with enduring deficits in behavioral inhibition in male mice,^65^ early adolescent treatment in females does not lead to changes in the proportion of commission errors incurred in the Go-No/Go task (Fig 2d). To obtained detailed information about individual performance across the No/Go task, we fit the proportion of commission error data of each mouse to a sigmoidal curve. Curve fitting revealed no differences in performance between treatment groups at the start of the task (upper asymptote, Fig 2e), in the number of days it took them to begin showing performance improvement (M50, Fig 2f), or in the proportion of commission errors made in the final trials (lower asymptote, Fig 2g). We also found no effect of AMPH in early adolescence on premature responses during training, a measure of waiting impulsivity (Extended data Fig 2b, c),^56,57^ nor in correct Go responses (Hits) during the task (Extended data Fig 2d).

**Table 2:**
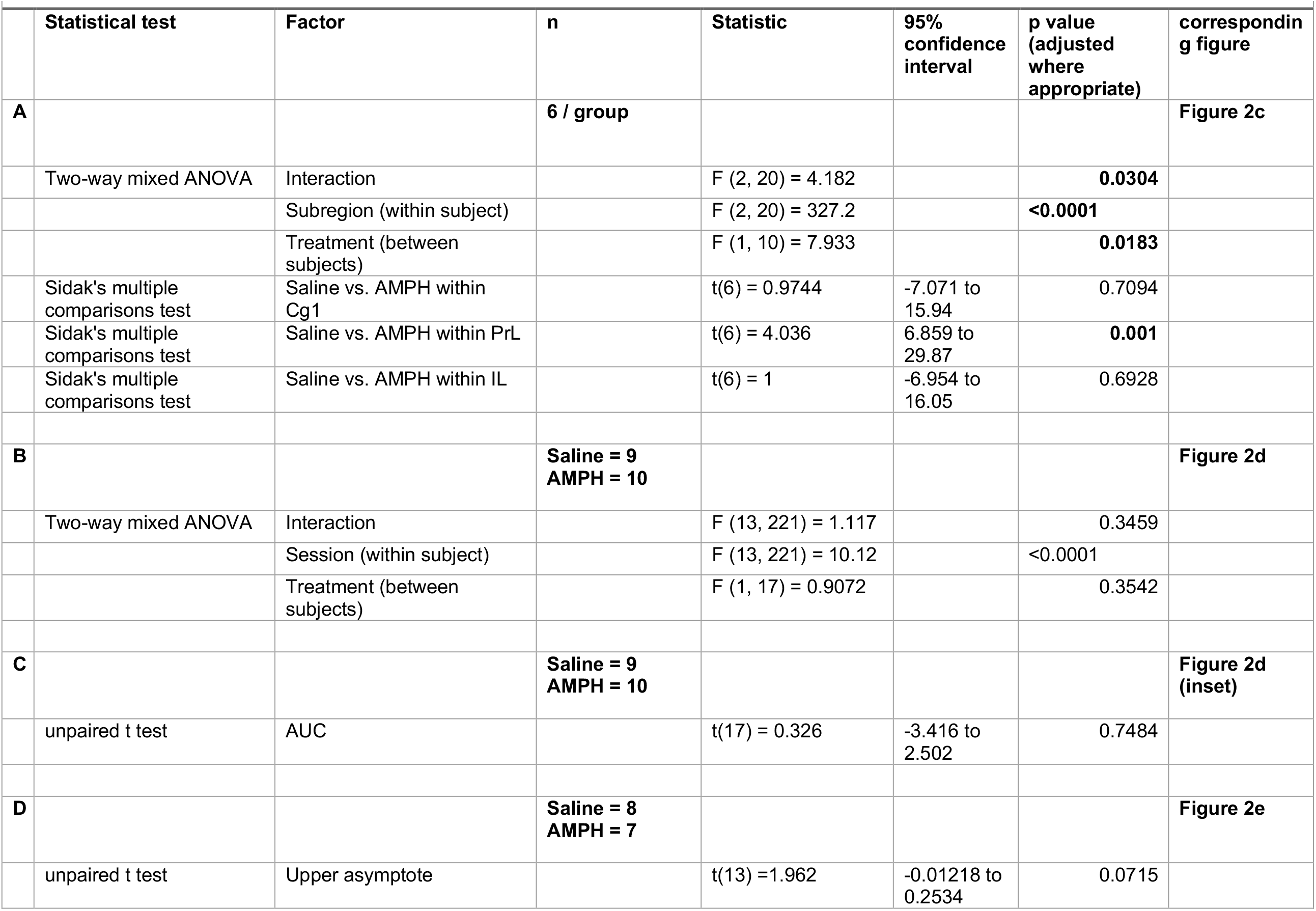
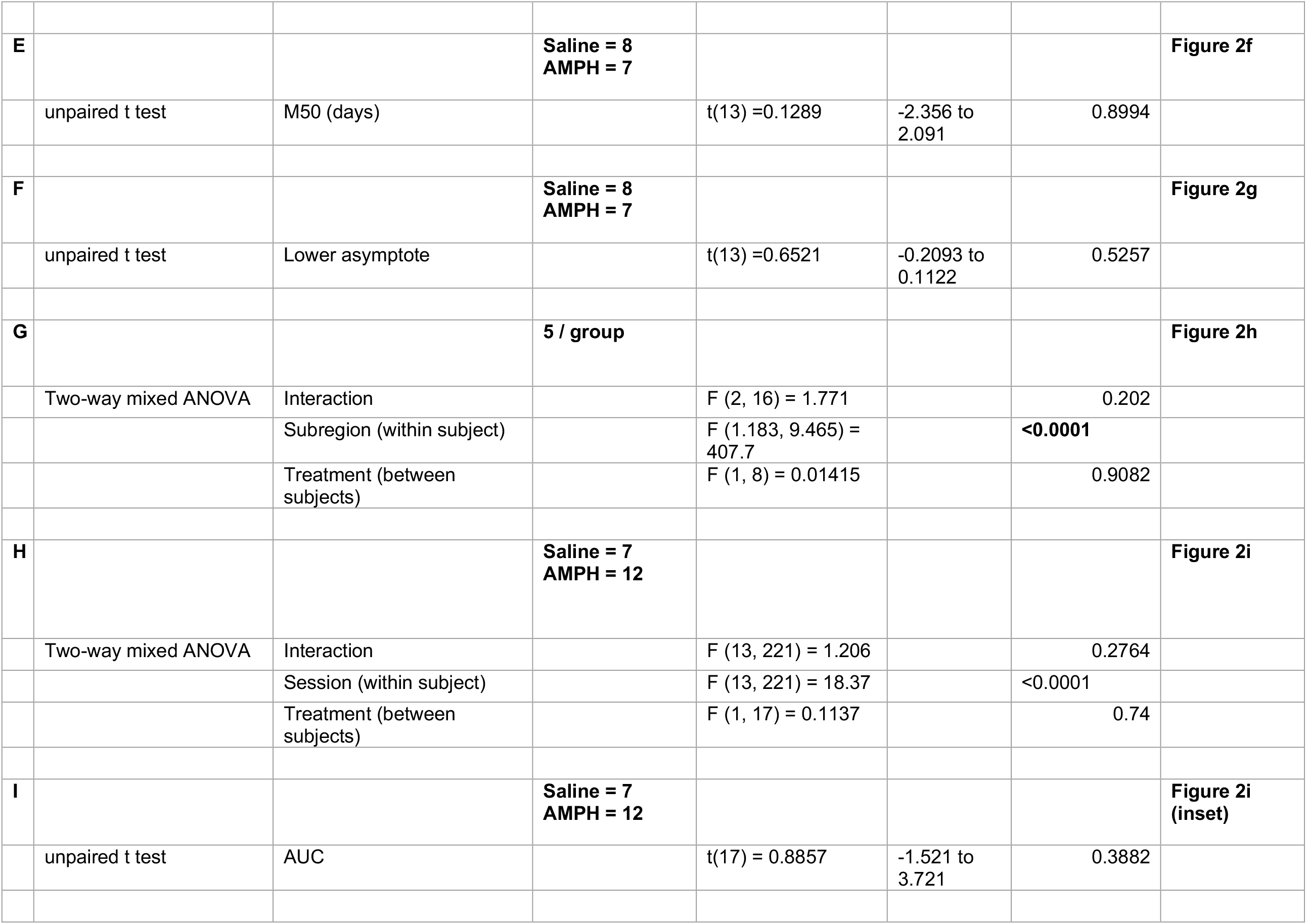
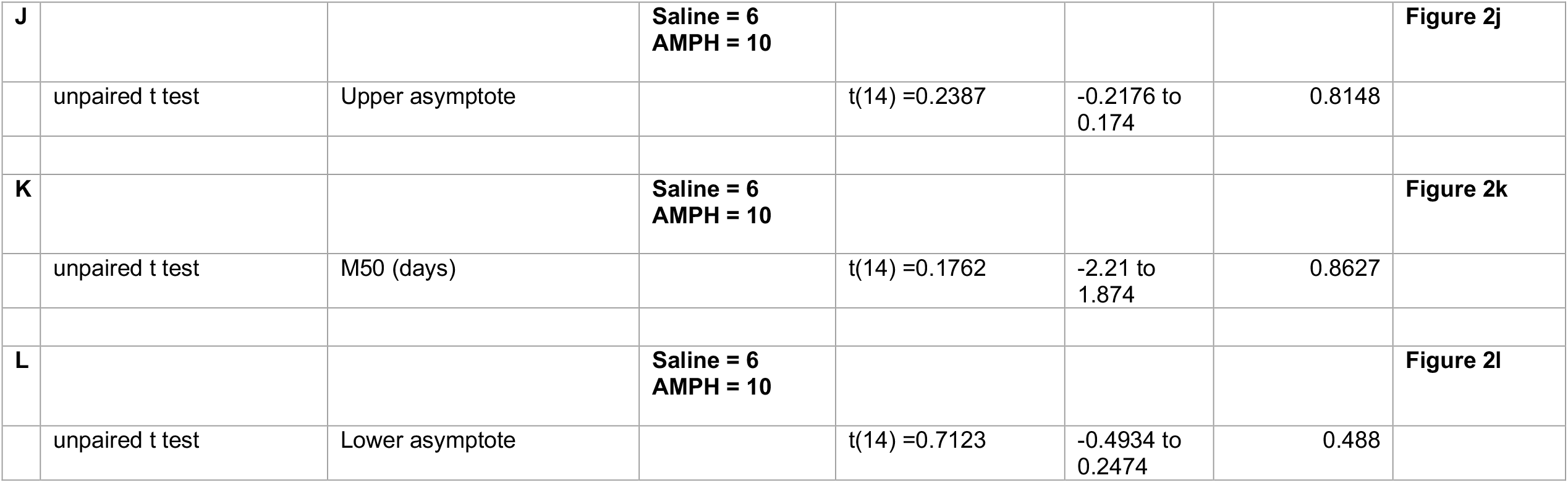
Detailed statistics for Figure 2.

Despite the ability of AMPH to downregulate *Dcc* levels in mid-adolescent female mice (Fig 1i), we found that exposure during this age period does *not* produce enduring changes in PFC dopamine innervation or in cognitive task performance (Fig 2h,i). Indeed, when fitting individual mouse performance to a sigmoidal curve, we found no differences between treatment groups in the upper asymptote (Fig 2j), M50 (Fig 2k), or lower asymptote (Fig 2l) measures. We also found no effect of AMPH in mid-adolescence on waiting impulsivity (Extended data Fig 2f, g), nor in correct Go responses (Hits) during the task (Extended data Fig 2h). These results suggest that sex- and age-specific compensatory processes may occur in the female mouse brain to counteract the downregulation of *Dcc* mRNA by AMPH in mid-adolescence.

### *Dcc* downregulation in dopamine neurons by AMPH in mid-adolescent females is compensated by Netrin-1 upregulation in the nucleus accumbens

The primary candidate for a compensatory process in mid-adolescent females is an opposing upregulation of Netrin-1, the ligand for DCC. Netrin-1 is expressed in the terminal regions of the mesocorticolimbic dopamine system, albeit in a complementary manner to the expression levels of DCC receptors in the dopamine axons that innervate these areas. In the PFC, Netrin-1 levels are high but only few dopamine axons express DCC. In contrast, Netrin-1 levels are lower in the nucleus accumbens (NAc), but all dopamine axons in this region highly express DCC (Fig 3a).^66^ Furthermore, dopamine axons are the only source of DCC protein expression in the NAc,^66^ suggesting a crucial and complementary role of DCC in dopamine axons and Netrin-1 in the NAc in the developmental organization of mesocorticolimbic dopamine connectivity. Finally, Netrin-1 levels in the NAc decline across adolescence in male mice,^61^ mirroring the same developmental pattern as we see in *Dcc* expression.^23–25^

**Figure 3.**
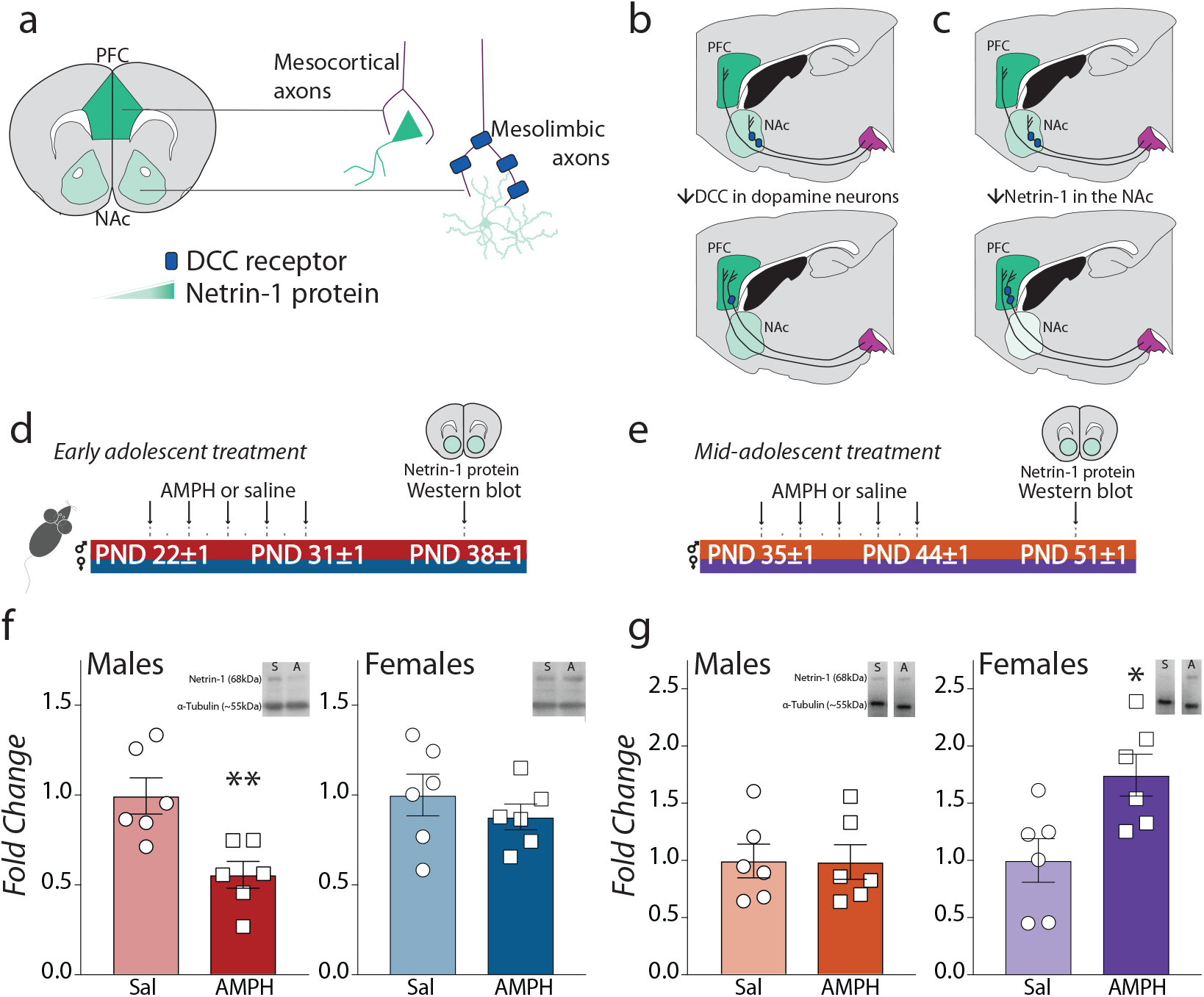
AMPH upregulates Netrin-1 in the NAc of mid-adolescent females, counteracting the downregulation it induces on Dcc levels in the VTA. *(a)* Expression of Netrin-1 in terminal regions of the mesocorticolimbic dopamine system. Netrin-1 is highly expressed in the PFC, with lower expression in the NAc. DCC is expressed in a complementary pattern, with DCC-expressing dopamine axons segregated to the NAc. *(b*) When *Dcc* is reduced in adolescent dopamine neurons using cre-lox recombination, axons with reduced *Dcc* fail to recognize the NAc as their final target and instead grow ectopically to the PFC.^53^ *(c)* Reducing Netrin-1 expression in the NAc with shRNA injection during adolescence also results in ectopic growth of *Dcc*-expressing dopamine axons to the PFC.^61^ *(d)* Timeline of experiments conducted in male and female mice treated with the recreational-like AMPH regimen in early adolescence (P21±1 to P31±1). AMPH in early adolescence downregulates NAc Netrin-1 levels in males (e), but not in females (f), compared to saline-treated controls (Table 3A,B). (g) Timeline of experiments conducted in mid-adolescent (P35±1 to P44±1) male and female mice treated with AMPH. While mid-adolescencent males are no longer sensitive to AMPH-induced downregulation of Netrin-1 in the NAc (h), mid-adolescent females show significant Netrin-1 *upregulation* in response to AMPH (i), opposing the drug-induced downregulation of *Dcc mRNA* in the VTA observed following this treatment (Figure 1i) (Table 3C, D). All graphs are normalized to the saline condition for each age and sex.

We thus next assessed the effects of AMPH treatment on Netrin-1 protein expression in the NAc of the same male and female mice in which we assessed *Dcc* mRNA levels in the VTA (Fig 3d, g; Extended data Fig 3). Of note, Netrin-1 is a ‘sticky’ guidance cue that accumulates on the surfaces of cells,^67–70^ thus quantification of protein levels gives the most functionally relevant account of its properties. We observed a significant reduction in Netrin-1 protein in the NAc of males, but not females, treated with AMPH in early adolescence in comparison to saline-treated controls (Fig 3e, f; Extended data Fig 3a). These findings are in line with our previous results in male mice,^58^ and show that the sex-specific effects of AMPH in early adolescence on *Dcc* expression in dopamine neurons extend to the regulation of Netrin-1 levels in the NAc.

**Table 3:**
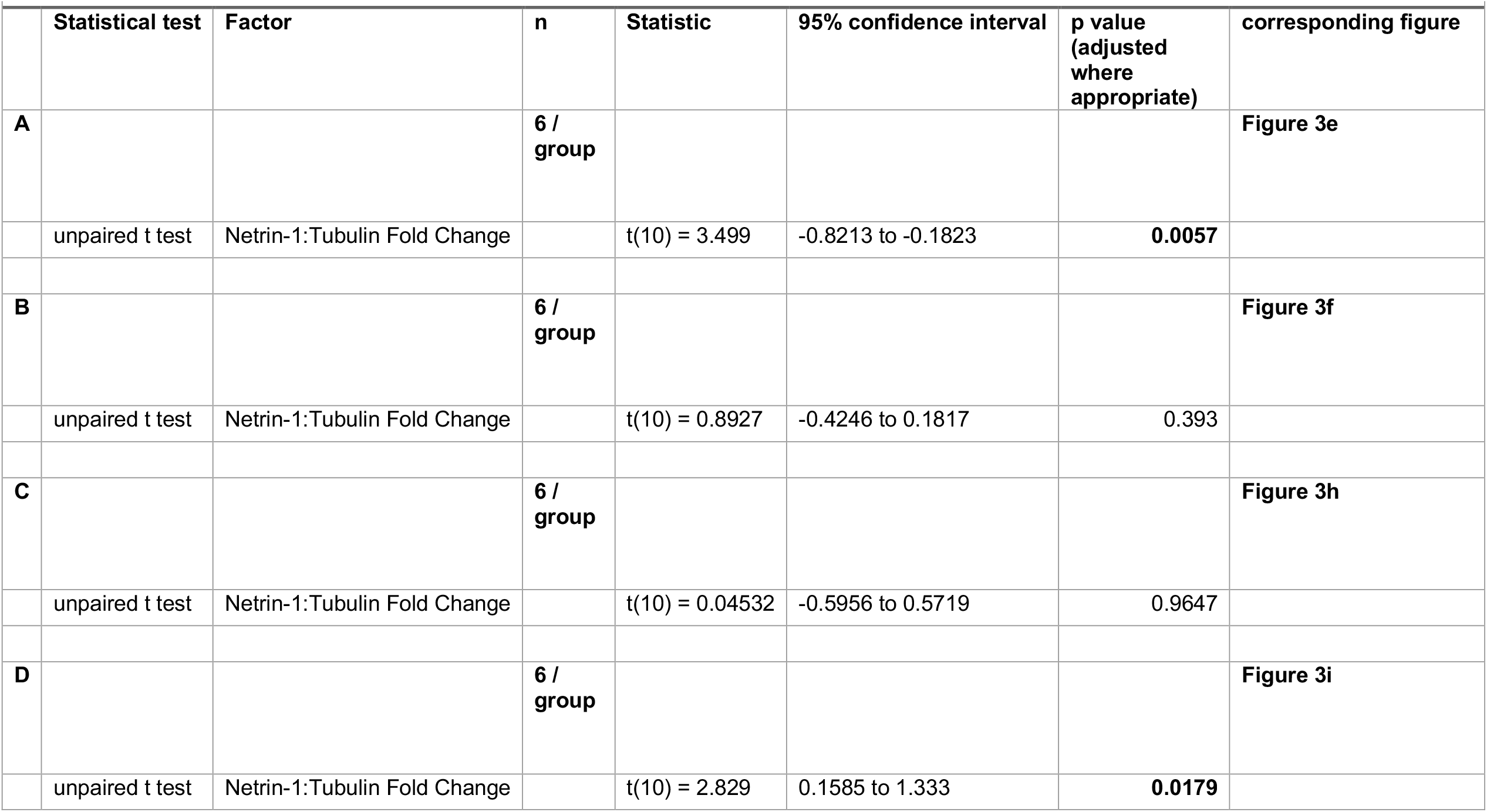
Detailed statistics for Figure 3.

|  | Statistical test | Factor | n | Statistic | 95% confidence interval | p value<br>(adjusted<br>where<br>appropriate) | corresponding figure |
| --- | --- | --- | --- | --- | --- | --- | --- |
| **A** |  |  | **6 /**<br>**group** |  |  |  | **Figure 3e** |
|  | unpaired t test | Netrin-1:Tubulin Fold Change |  | t(10) = 3.499 | -0.8213 to -0.1823 | **0.0057** |  |
| **B** |  |  | **6 /**<br>**group** |  |  |  | **Figure 3f** |
|  | unpaired t test | Netrin-1:Tubulin Fold Change |  | t(10) = 0.8927 | -0.4246 to 0.1817 | 0.393 |  |
| **C** |  |  | **6 /**<br>**group** |  |  |  | **Figure 3h** |
|  | unpaired t test | Netrin-1:Tubulin Fold Change |  | t(10) = 0.04532 | -0.5956 to 0.5719 | 0.9647 |  |
| **D** |  |  | **6 /**<br>**group** |  |  |  | **Figure 3i** |
|  | unpaired t test | Netrin-1:Tubulin Fold Change |  | t(10) = 2.829 | 0.1585 to 1.333 | **0.0179** |  |

Exposure to AMPH in mid-adolescence (Figure 3g) does not alter Netrin-1 in the NAc of male mice (Fig 3h, Extended data Fig 3b), consistent with their lack of sensitivity to the later timing of this drug treatment. Notably, however, AMPH in mid-adolescent females significantly *upregulates* Netrin-1 protein expression in the NAc (Fig 3i, Extended data Fig 3b), indicating that at this later adolescent age, AMPH induces *opposite* regulation of DCC receptors in dopamine neurons and of Netrin-1 in their mesolimbic targets. Upregulation of Netrin-1 in females may be a compensatory effect of drug treatment, protecting against the enduring consequences triggered by drug-induced *Dcc* downregulation.

### AMPH in early adolescence induces ectopic growth of mesolimbic dopamine axons to the PFC in male mice

How exactly AMPH exposure in early adolescence produces enduring changes to PFC dopamine structure and cognitive function in male mice has remained a matter of debate. In contrast to the results in female mice (Fig 2c, h), we found an increase in the span of the dopamine input to the PFC in adult males exposed to AMPH in early adolescence (Extended data Fig 4a). To determine the origin of this increase, we used intersectional viral tracing (Fig 4a)^53^ in male mice exposed to AMPH or to saline in early adolescence, to track the growth of dopamine axons in adolescence as they make targeting decisions at the level of the NAc. We looked specifically at the eYFP+ axons, which represent dopamine axons labeled at the level of the NAc at the start of adolescence. We found significantly more eYFP+ dopamine axon terminals in the PFC of adult mice that were exposed to AMPH in early adolescence, in comparison to saline-exposed counterparts (Fig 4b). This effect was more pronounced in ventral PFC subregions (Fig 4c), in line with the overall changes in dopamine input volume seen in the same brain sections (Extended data Fig 4a). In addition, the number of eYFP+ dopamine terminals in the NAc is reduced in AMPH groups compared to saline controls (Fig 4d), and there is a strong negative correlation between PFC and NAc eYFP+ dopamine terminals (Fig 4e), indicating that AMPH in adolescence reroutes dopamine axons intended to innervate the NAc to the PFC. Minor changes in dopamine innervation to the PFC in adolescence can substantially shape the morphology of postsynaptic neurons.^53,55^ Indeed we find significant restructuring of pyramidal neuron arbors and changes in their spine density after AMPH in early adolescence (Extended data Fig 4 c-g).

**Figure 4.**
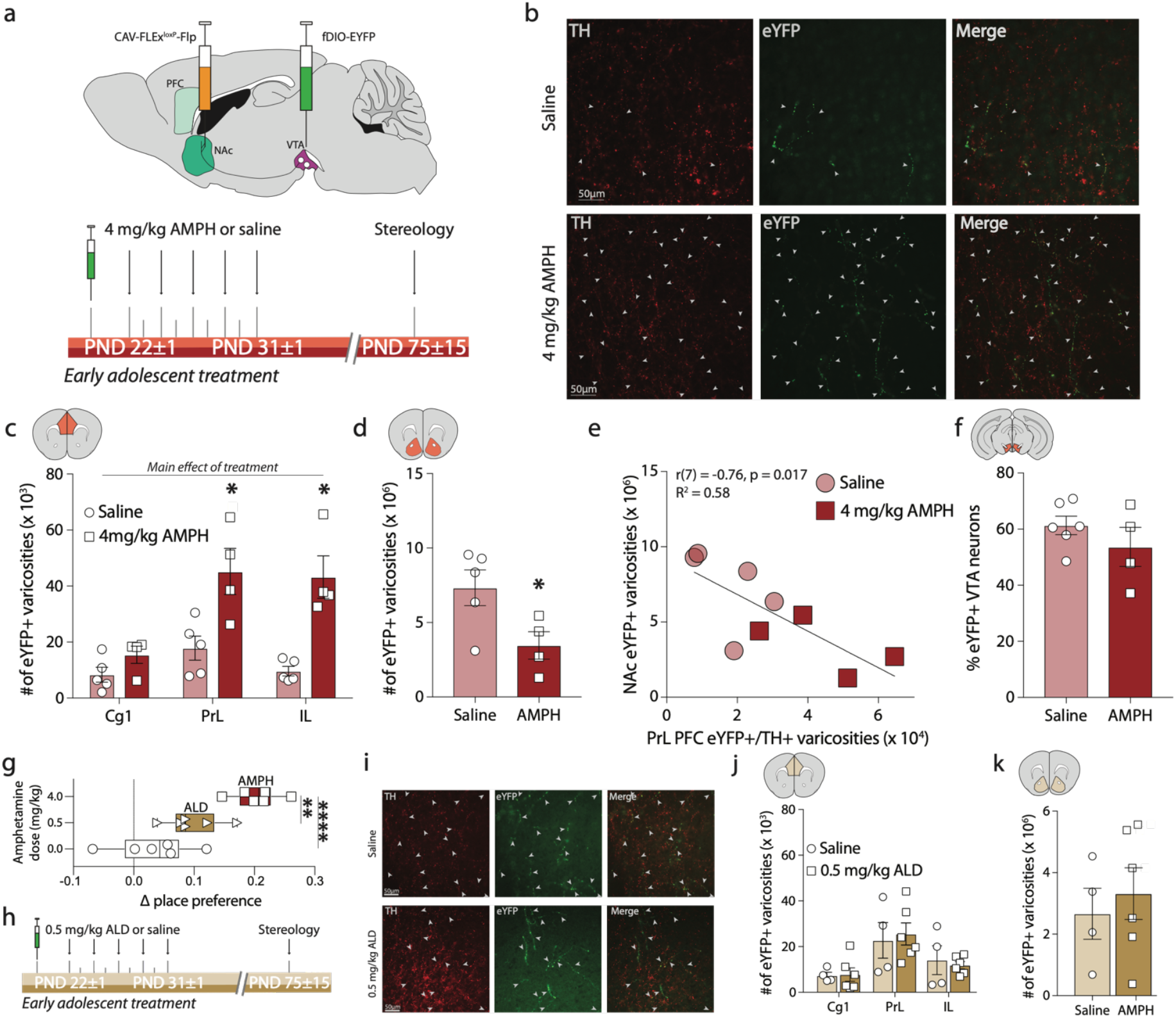
Recreational AMPH in adolescence induces ectopic growth of mesolimbic dopamine axons to the PFC in male mice. (a) Experimental design. Tracer viruses were injected in *DAT*^*Cre*^ mice at PND 21 to limit recombination only to dopamine neurons with axons terminals in the NAc at that age. Mice were then exposed to the recreational-like amphetamine (AMPH, 4 mg/kg) regimen from P21±1 to P31±1,^62^ a treatment consistently shown to decrease DCC receptor expression in the VTA in early adolescence, to disrupt PFC dopamine connectivity, and to induce deficits in impulse control in adulthood.^58,64,65,88^ (b) Photomicrographs of coronal sections showing the prelimbic PFC (PrL) of adult mice injected with tracer viruses in adolescence. *Top* Mice injected with saline during adolescence show that dopamine axons continued to grow from the NAc to the PFC in adolescence (closed arrowheads), consistent with our previous results in drug-naive mice.^53^ *Bottom* The number of axons that grew to the PFC in adolescence is dramatically increased in adult mice that were exposed to AMPH early in adolescence. (c) Stereological quantification reveals a significant increase in fluorescent axon terminals across the cingulate 1 (Cg1), prelimbic PrL, and infralimbic (IL) subregions of the medial PFC and a pronounced dorsal-to-ventral gradient (Table 4A). (d) The number of fluorescently labeled terminals in the NAc is significantly decreased in AMPH -treated mice in comparison to those that received saline (Table 4B). (e) The number of labeled terminals in the PFC and in the NAc are negatively correlated (Table 4C). (f) The percentage of VTA dopamine neurons infected by the tracing viruses is similar between treatment groups (Table 4D). (g) Exposure to AMPH induces robust conditioned place preference (CPP) (Table 1E). This is not the case in male mice exposed to a therapeutic-like amphetamine regimen (ALD, 0.5 mg/kg). (h) Experimental design. Tracer viruses were injected in *DAT*^*Cre*^ mice as in (*d*), but mice were then exposed to ALD, which does *not* alter *Dcc* mRNA levels in the VTA in adolescence.^62^ (i) Photomicrographs of coronal sections showing the PrL of adult mice injected with tracer viruses in adolescence. The number of labeled axons that continued to grow from the NAc to the PFC in adolescence (closed arrowheads) is not different between adult mice that were exposed to saline (*Top*) or to ALD in adolescence (*Bottom*). (j-k) ALD does not alter dopamine axon growth to the Cg1, PrL, or IL in adolescence (j, Table 1F) or the number of labeled dopamine terminals in the NAc (k, Table 1G), indicating that this AMPH dose does not interfere with dopamine axon targeting.

To determine whether the ectopic growth of mesolimbic dopamine axons to the PFC is induced specifically by recreational-like doses of amphetamine (AMPH) in early adolescent male mice, we investigated whether an Adderall-like dose (0.5 mg/kg d-amphetamine, ALD), which produces similar plasma levels in mice as therapeutic treatment with d-amphetamine (trade name Adderall) in humans,^62^ would produce similar effects on PFC dopamine development. While non-contingent AMPH induces a robust place preference in adolescent mice, the same treatment regimen with an ALD does not (Figure 4g). This is in agreement with earlier studies indicating that ALD do not alter dopamine system function in rats,^71^ and do not induce long-term changes to PFC function, including deficits in inhibitory control in rodents or in non-human primates.^62,72^ To investigate if the rerouting effect of AMPH on NAc dopamine axons is dose-dependent, we performed the same experiments as in Figure 4a, but comparing saline versus ALD administration (Figure 4h). ALD in early adolescence does not alter dopamine axon growth to the PFC (Figure 4i, j), does not produce changes in the volume of dopamine innervation (Extended data Fig 4b), nor does it change the number of eYFP+ dopamine terminals remaining in the NAc (Figure 4k). The disruptive effects of amphetamine on dopamine development are linked to properties specific to recreational-like AMPH doses.

**Table 4:**
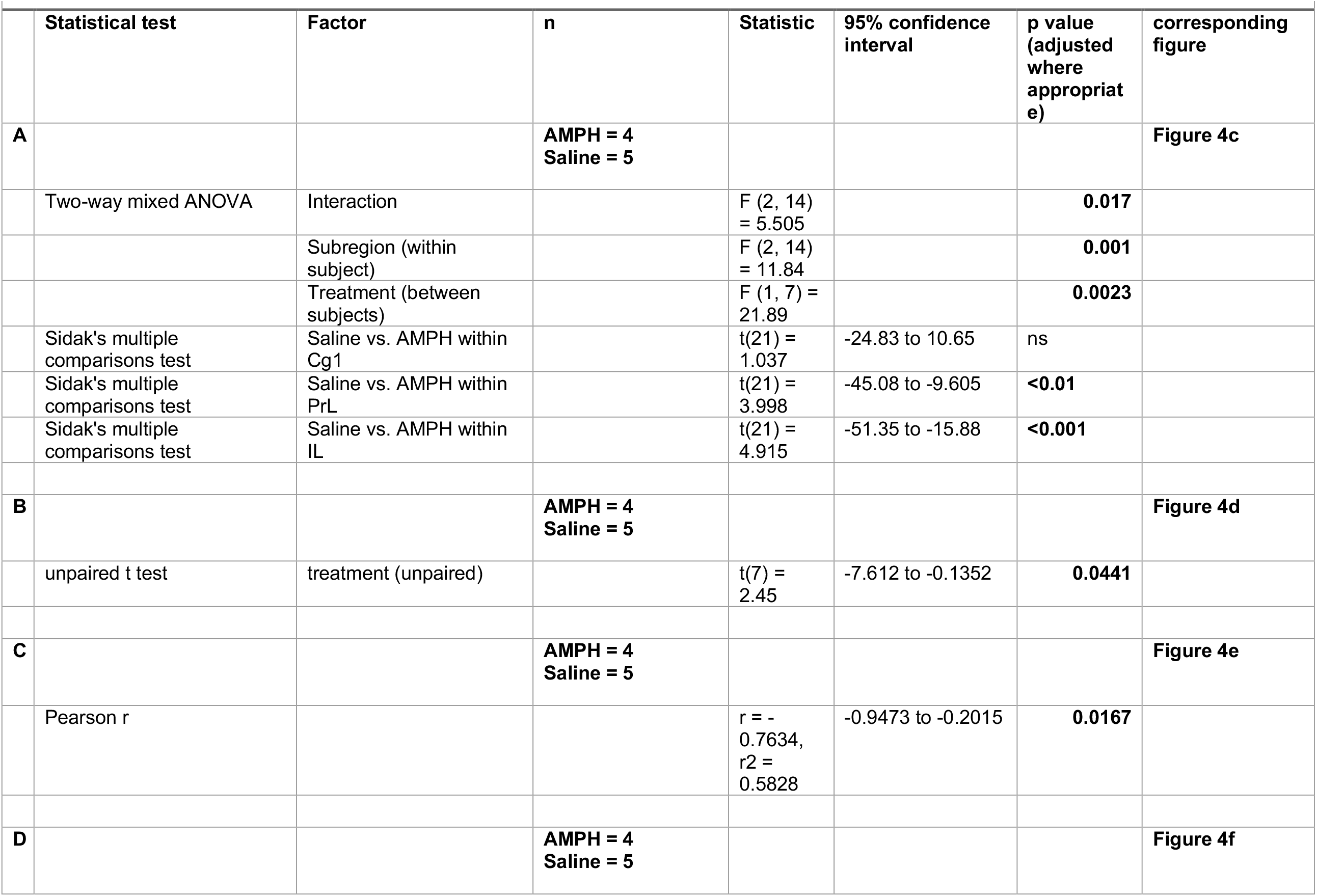
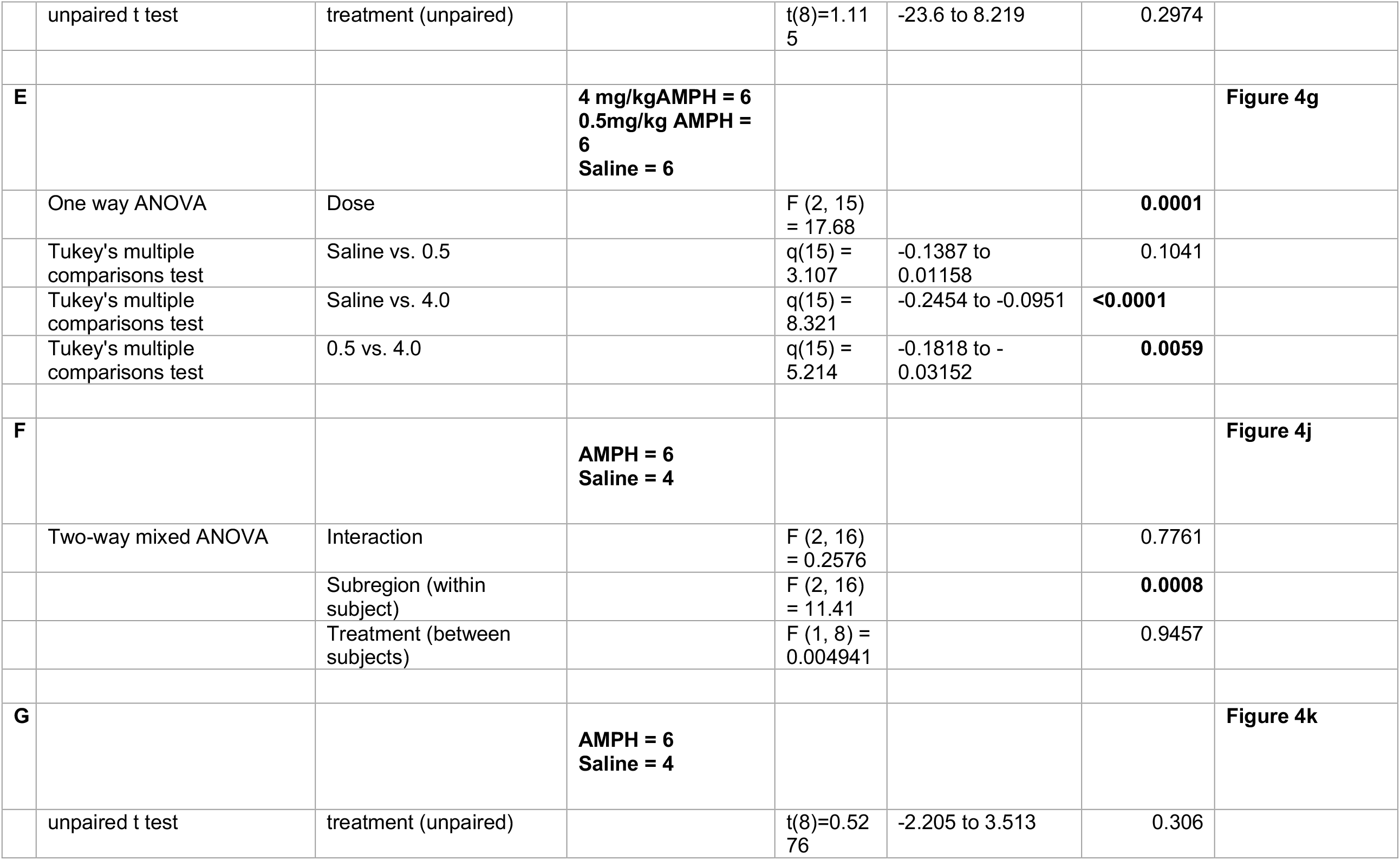
Detailed statistics for Figure 4.

| Statistical test | Factor | n | Statistic | 95% confidence interval | p value (adjusted where appropriate) | corresponding figure |
| --- | --- | --- | --- | --- | --- | --- |
| **A** |  | **AMPH = 4**<br>**Saline = 5** |  |  |  | **Figure 4c** |
| Two-way mixed ANOVA | Interaction |  | F (2, 14) = 5.505 |  | **0.017** |  |
|  | Subregion (within subject) |  | F (2, 14) = 11.84 |  | **0.001** |  |
|  | Treatment (between subjects) |  | F (1, 7) = 21.89 |  | **0.0023** |  |
| Sidak's multiple comparisons test | Saline vs. AMPH within Cg1 |  | t(21) = 1.037 | -24.83 to 10.65 | ns |  |
| Sidak's multiple comparisons test | Saline vs. AMPH within PrL |  | t(21) = 3.998 | -45.08 to -9.605 | **<0.01** |  |
| Sidak's multiple comparisons test | Saline vs. AMPH within IL |  | t(21) = 4.915 | -51.35 to -15.88 | **<0.001** |  |
| **B** |  | **AMPH = 4**<br>**Saline = 5** |  |  |  | **Figure 4d** |
| unpaired t test | treatment (unpaired) |  | t(7) = 2.45 | -7.612 to -0.1352 | **0.0441** |  |
| **C** |  | **AMPH = 4**<br>**Saline = 5** |  |  |  | **Figure 4e** |
| Pearson r |  |  | r = -0.7634,<br>r<sup>2</sup> = 0.5828 | -0.9473 to -0.2015 | **0.0167** |  |
| **D** |  | **AMPH = 4**<br>**Saline = 5** |  |  |  | **Figure 4f** |

|  |  |  |  |  |  |  |  |
| --- | --- | --- | --- | --- | --- | --- | --- |
| | unpaired t test | treatment (unpaired) | | $t(8)=1.11$<br>5 | -23.6 to 8.219 | 0.2974 | |
| **E** |  |  | **4 mg/kg AMPH = 6**<br>**0.5mg/kg AMPH = 6**<br>**Saline = 6** |  |  |  | **Figure 4g** |
| | One way ANOVA | Dose | | $F(2, 15) = 17.68$ | | **0.0001** | |
| | Tukey's multiple comparisons test | Saline vs. 0.5 | | $q(15) = 3.107$ | -0.1387 to 0.01158 | 0.1041 | |
| | Tukey's multiple comparisons test | Saline vs. 4.0 | | $q(15) = 8.321$ | -0.2454 to -0.0951 | **<0.0001** | |
| | Tukey's multiple comparisons test | 0.5 vs. 4.0 | | $q(15) = 5.214$ | -0.1818 to -0.03152 | **0.0059** | |
| **F** |  |  | **AMPH = 6**<br>**Saline = 4** |  |  |  | **Figure 4j** |
| | Two-way mixed ANOVA | Interaction | | $F(2, 16) = 0.2576$ | | 0.7761 | |
| | | Subregion (within subject) | | $F(2, 16) = 11.41$ | | **0.0008** | |
| | | Treatment (between subjects) | | $F(1, 8) = 0.004941$ | | 0.9457 | |
| **G** |  |  | **AMPH = 6**<br>**Saline = 4** |  |  |  | **Figure 4k** |
| | unpaired t test | treatment (unpaired) | | $t(8)=0.52$<br>76 | -2.205 to 3.513 | 0.306 | |

### A *Dcc*-dependent mechanism underlies the enduring deficits in impulse control induced by AMPH exposure in early adolescence

Exposure to AMPH, but not ALD, early in adolescence decreases *Dcc* mRNA expression in the VTA of male mice,^62^ where 99% of dopamine neurons express *Dcc*.^59^ Our previous work indicates that *Dcc* expression in the VTA is important for appropriate dopamine axon targeting in adolescence, when the mesocorticolimbic dopamine pathway is still actively developing,^53^ suggesting DCC receptors in dopamine axons as mediators of the effects of AMPH effects on mesocorticolimbic dopamine development.^64^ However, a causal relationship between AMPH-induced changes in *Dcc* expression and impulsivity has never been addressed, owing to limitations in tools to manipulate *Dcc* levels. While germ line and conditional knock-downs of *Dcc* expression have been investigated by our team and others, interventions to *increase Dcc* expression levels has been challenging to achieve due in part to the large size of the *Dcc* gene and mRNA, as *Dcc* cDNA is too large to be packaged in traditional AAV over-expression vectors (Fig 5a).^73,74^ To be able to establish if *Dcc* mediates the effects of AMPH in adolescence on dopamine axon rerouting and on enduring cognitive impairments, we designed a CRISPR activation (CRISPRa) system to specifically upregulate the transcription of the *Dcc* gene in mice (Fig 5b)^75^. In vitro characterization (Fig 5c) using dopamine cell cultures shows a moderate increase in *Dcc* mRNA expression from each of the 4 single guide RNA (sgRNA) sequences tested when compared to the *LacZ* sgRNA control (Fig 5d). Single gene multiplexing (Fig 5e) – i.e. combinatorial application of sgRNAs – indicates that a nearly 4-fold increase in *Dcc* mRNA could be achieved by combining the 4 sequences. For all of the following experiments, this cocktail of the 4 *Dcc* sgRNAs was used. In vivo testing (Fig 5f) revealed that the sgRNAs can be well expressed in the VTA of adolescent male mice (Fig 5g), and that sgRNA expression is observed in VTA dopamine neurons (Fig 5h). Quantitative analyses revealed a significant upregulation of *Dcc* mRNA expression in the VTA (Fig 5i), as well as a significant increase in DCC protein expression in the NAc (Fig 5j), where DCC protein is *not* expressed by local cells – it localizes *only* to dopamine axons.^66^ The expression of DCC protein in NAc dopamine axons is strongly correlated with mRNA expression in the VTA (Fig 5k), indicating that the CRISPRa system produces robust upregulation of *Dcc* mRNA transcription, ultimately increasing protein translation and localization throughout the neuron.

**Figure 5.**
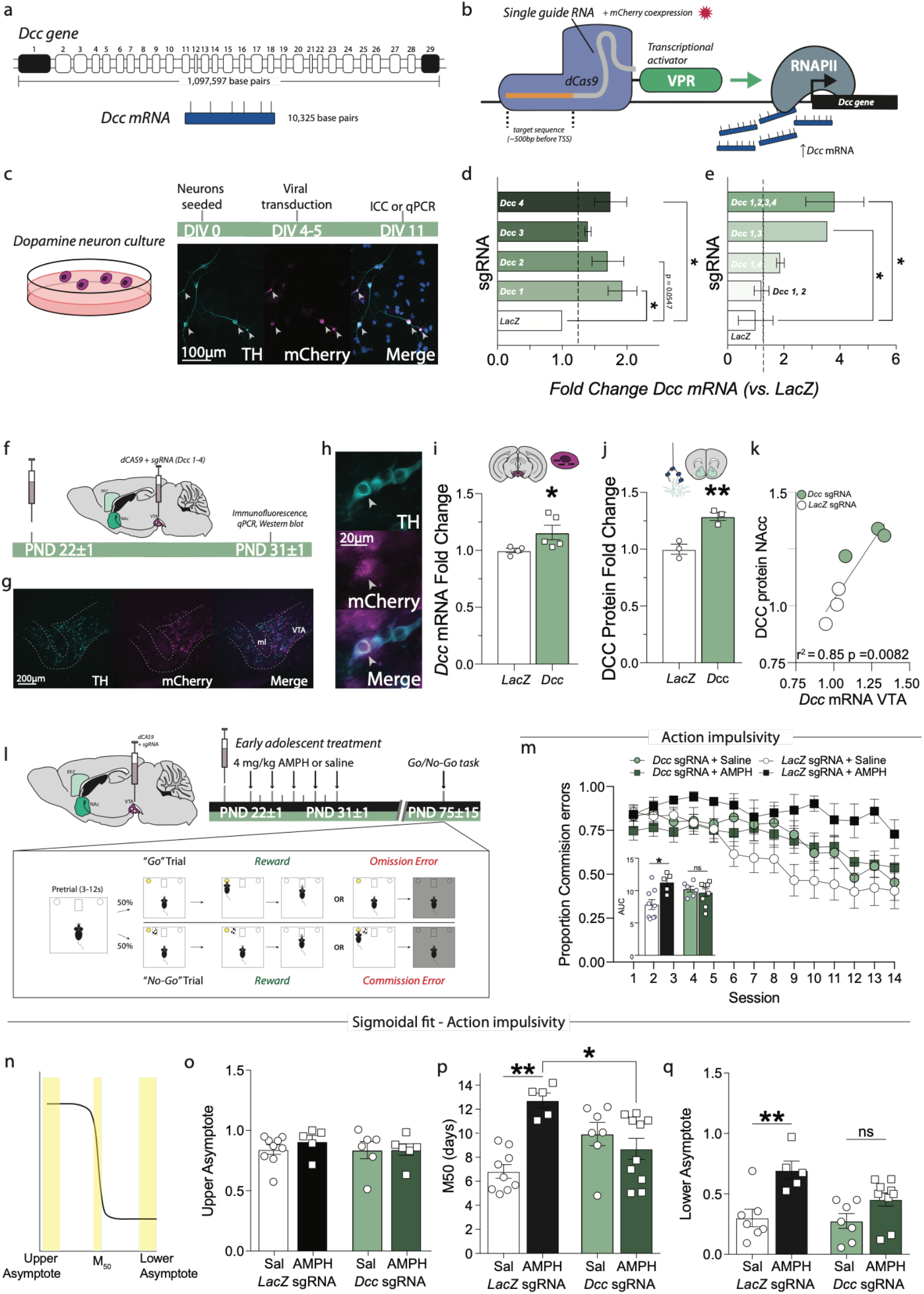
Upregulation of Dcc transcription in the VTA with CRISPRa prevents the harmful effect of recreational AMPH in adolescence on impulse control. (a) The murine *Dcc* gene spans 29 exons, contains more than 1 million base pairs,^73,89,90^ and encodes an mRNA of over 10 kilobases (NCBI reference sequence: NM_007831.3). Because of its large size, *Dcc* is not readily amenable to typical cDNA overexpression approaches. (b) CRISPR activation (CRISPRa) system.^75^ (c) Dopamine cell cultures were used to validate expression of sgRNA sequences in transfected cells. Photomicrograph shows cultured dopamine neurons at DIV 11 expressing the mCherry tag for *Dcc* sgRNA, as revealed by co-immunofluorescence of mCherry and TH (arrowheads). (d) Four sgRNA sequences targeting different regions ∼500bp upstream of the transcription start site (TSS) of the *Dcc* gene were tested in dopamine neuron cultures, using an sgRNA targeting *LacZ* as a control. While all sequences augmented *Dcc* mRNA expression (Table 5A; minimum 1.3983± 0.04752 fold change), the increase was limited (maximum 1.93135± 0.22296 fold change). (e) A single gene multiplexing procedure was used to maximize the increase in *Dcc* mRNA overexpression by pooling the 4 *Dcc* sgRNA sequences in the same injection. The multiplex of all 4 sgRNAs gave the most robust increase in *Dcc* mRNA, with a maximal fold change of 3.81668 ±1.03421 over *LacZ* control (Table 5B). (f) In vivo experimental design. Lentiviruses expressing dCas9-VPR and the 4 *Dcc* sgRNAs (or the *LacZ* control) were injected bilaterally into the VTA of male mice at PND 21. On PND 31 brains were harvested and processed for immunofluorescence or for qPCR and Western blot. (g) Low and (h) high magnification image of tyrosine hydroxylase (TH)+ dopamine neurons expressing the sgRNA viruses (mCherry; arrowhead). (i) qPCR revealed a significant upregulation of *Dcc* mRNA in the VTA of mice 10 days after receiving *Dcc* sgRNAs compared to those receiving the *LacZ* sgRNA (Table 5C). (j) Notably, DCC protein expression was significantly increased in the NAc, where DCC receptors are present only on dopamine axons (Table 5D). (k) Indeed, DCC protein upregulation in the NAc was strongly correlated with *Dcc* mRNA upregulation in the VTA (Table 5E). (l) *Top* Experimental design. Male mice were injected with viruses expressing dCas9-VPR and the 4 *Dcc* sgRNAs multiplex as in (*e*). Following viral microinfusions, mice were exposed to the recreational-like AMPH regimen.^62^ Mice were then allowed to grow to adulthood, when they were tested in an operant Go/No-Go task. *Bottom* During the Go/No-Go task mice have to respond to a light cue to receive a food reward, but to withhold their response when a tone is paired with the light cue. (m) Action impulsivity was measured by calculating the proportion of commission errors during the No-Go part of the task. Mice with *LacZ* sgRNA microinfusions and treated with AMPH in adolescence showed a greater rate of commission errors throughout the 14-day task compared to mice with *LacZ* sgRNA and treated with saline, indicating greater action impulsivity or impaired behavioral inhibition. This effect of AMPH was not observed in mice that received *Dcc* sgRNA (Table 5G). Area under the curve (AUC, inset) indicates that *Dcc* CRISPRa protects against AMPH-induced action impulsivity (Table 5H). (n) Illustration of sigmoidal curve fit analysis. (o) Mice across all groups showed similar number of commission errors at the beginning of the task (upper asymptote, Table 5I). (p) *LacZ* sgRNA AMPH-treated mice took longer to improve their task performance (M50) in comparison to the *LacZ* sgRNA saline group (Table 5J), with some mice never improving (an M50 of 14 days). In contrast, *Dcc* CRISPRa mice treated with AMPH in adolescence did not take longer to improve in the task than the *Dcc* CRISPRa mice treated with saline and show performance improvement significantly earlier than the AMPH-treated LacZ sgRNA mice. (q) During the last trials, only *LacZ* sgRNA AMPH-treated mice showed significant impulse control deficits (Table 2K).

We next asked if restoring *Dcc* expression in dopamine neurons with CRISPRa could block the effects of AMPH in early adolescence on deficits in behavioral inhibition in adulthood.^65^ Male mice were injected with the sgRNA cocktail and dCas9 viruses at PND21 and then treated with a regimen of saline or AMPH (Fig 5l). All mice were then tested in a Go-No/Go task in adulthood. All adult mice treated with saline in adolescence showed a marked reduction in commission errors across the 14 testing days, whether they received CRISPRa for *Dcc* (*Dcc* sgRNA) or the *LacZ* sgRNA control, indicative in an improvement in action inhibition across the task (Figure 5m). Adult mice that received CRISPRa with *LacZ* sgRNA and were treated with AMPH in early in adolescence show deficits in behavioral inhibition, in line with previous results.^65^ Strikingly, receiving CRISPRa with the *Dcc* sgRNAs prevents the development of persistent action impulsivity induced by AMPH exposure in adolescence, as this mouse group does not perform significantly different than the saline-treated groups (Fig 5m, *inset*). To determine when in the task the differences in performance across groups emerged, individual task performance data were fitted to a sigmoidal curve (Fig 5n). The resulting analysis reveals that while there are no differences in initial performance (Fig 5o, *upper asymptote*), mice that received CRISPRa with the *LacZ* sgRNA and were treated with AMPH in early adolescence take significantly longer to show improvement in the No/Go task than the other groups, evidenced by a greater M_50_ (Fig 5p). Notably, some AMPH-treated control mice do not improve at all over the task, since their M_50_ value is equal to the total number of sessions. In contrast, there is no difference in the M_50_ between the saline and AMPH treated mice that received CRISPRa with the *Dcc* sgRNA, indicative that CRISPRa-mediated *Dcc* upregulation protects against the enduring effect of AMPH on action inhibition. Finally, while AMPH treatment in early adolescence significantly increases the value of the lower asymptote in *LacZ* sgRNA groups, indicating profound impulse control deficits (Fig 5q), this effect is blocked by the CRISPRa treatment targeting *Dcc* in the VTA.

**Table 5:**
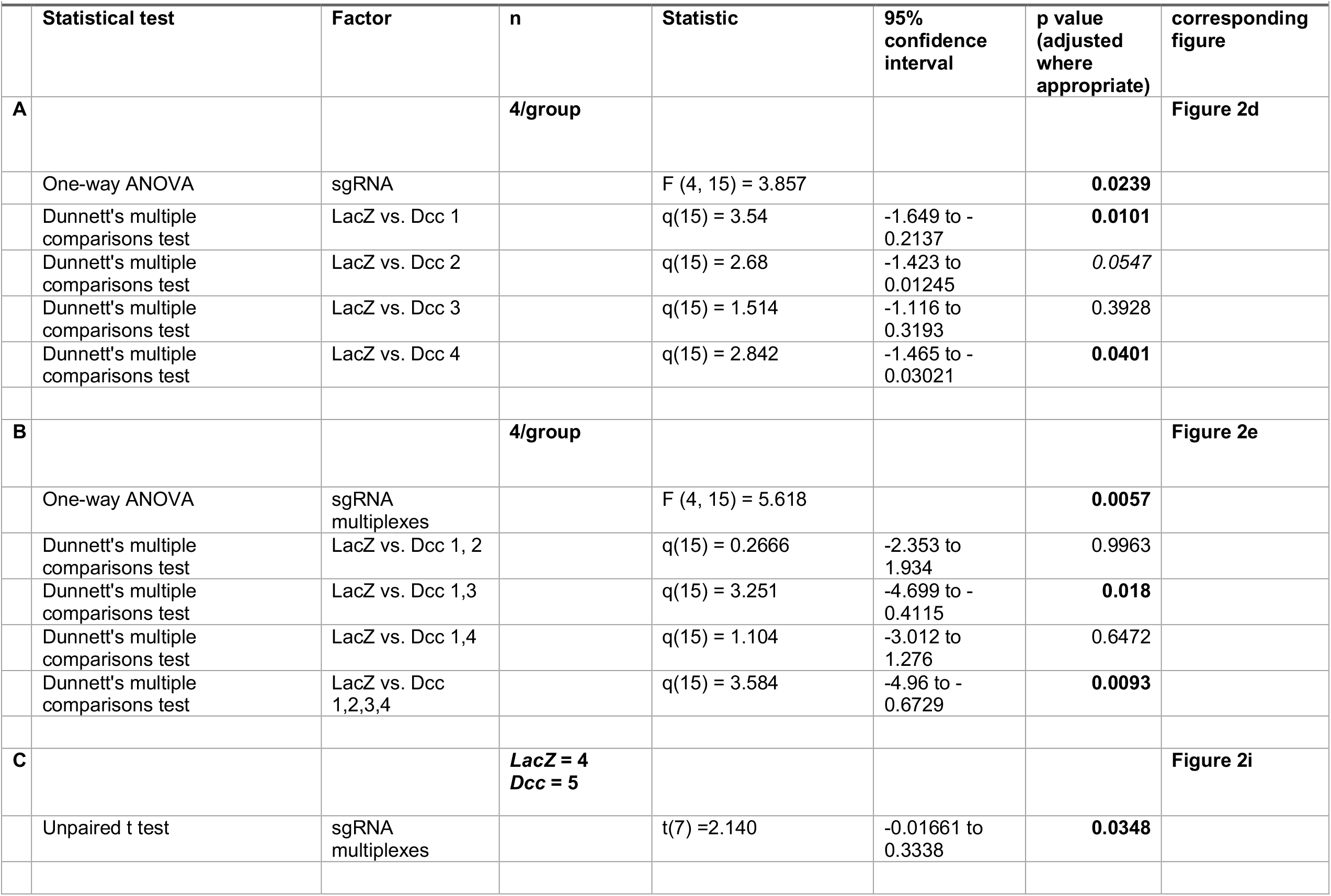
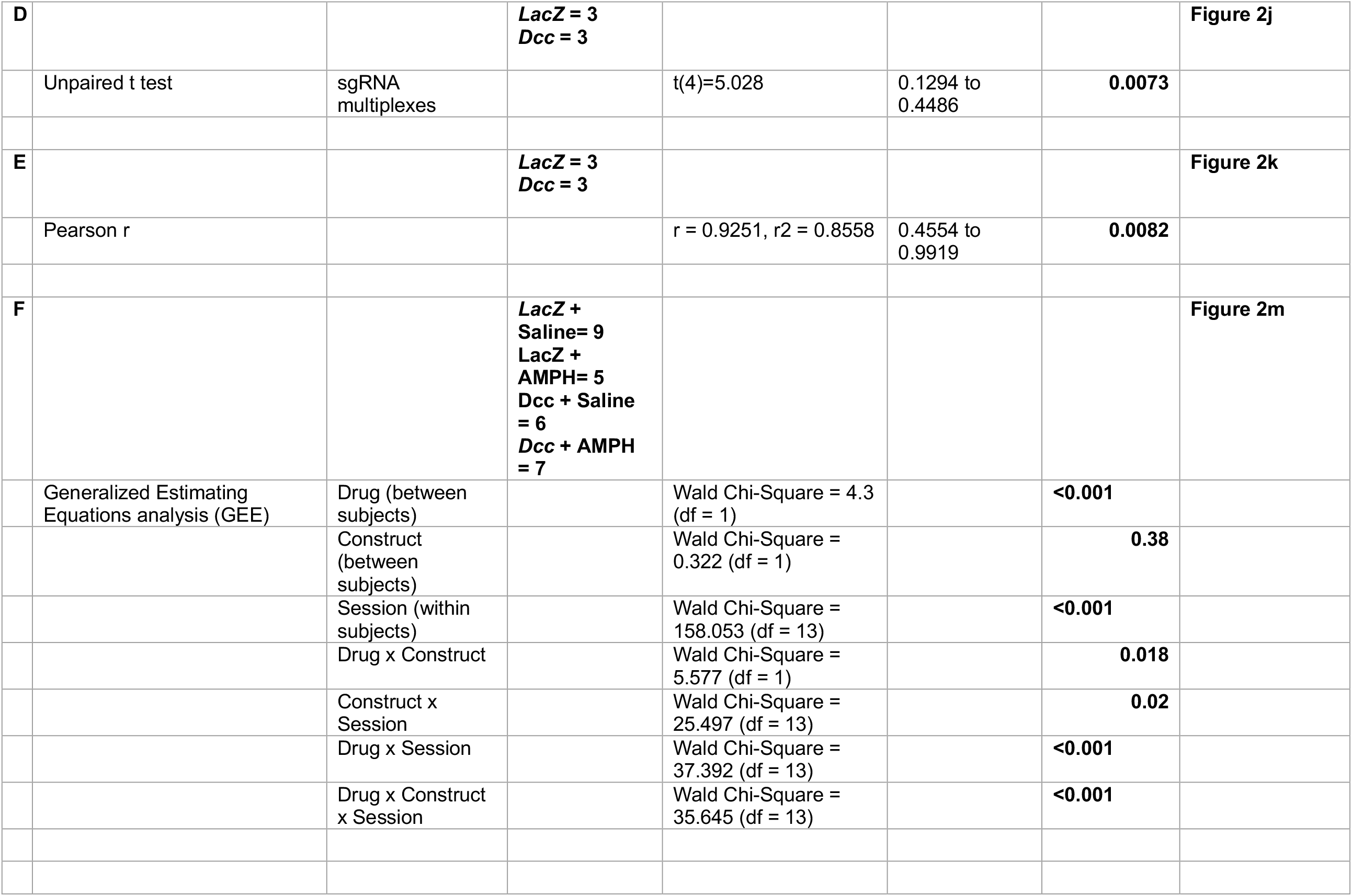
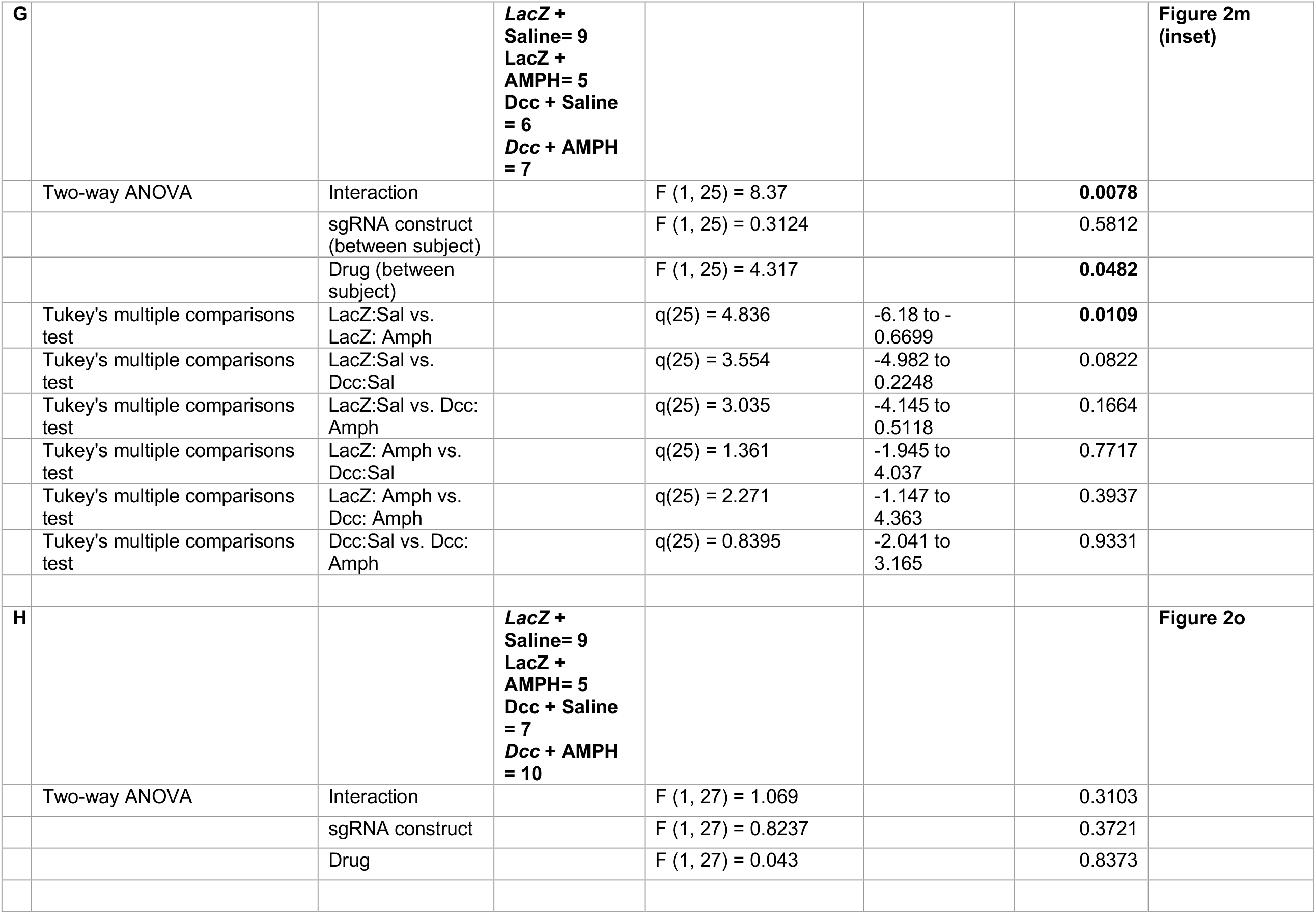
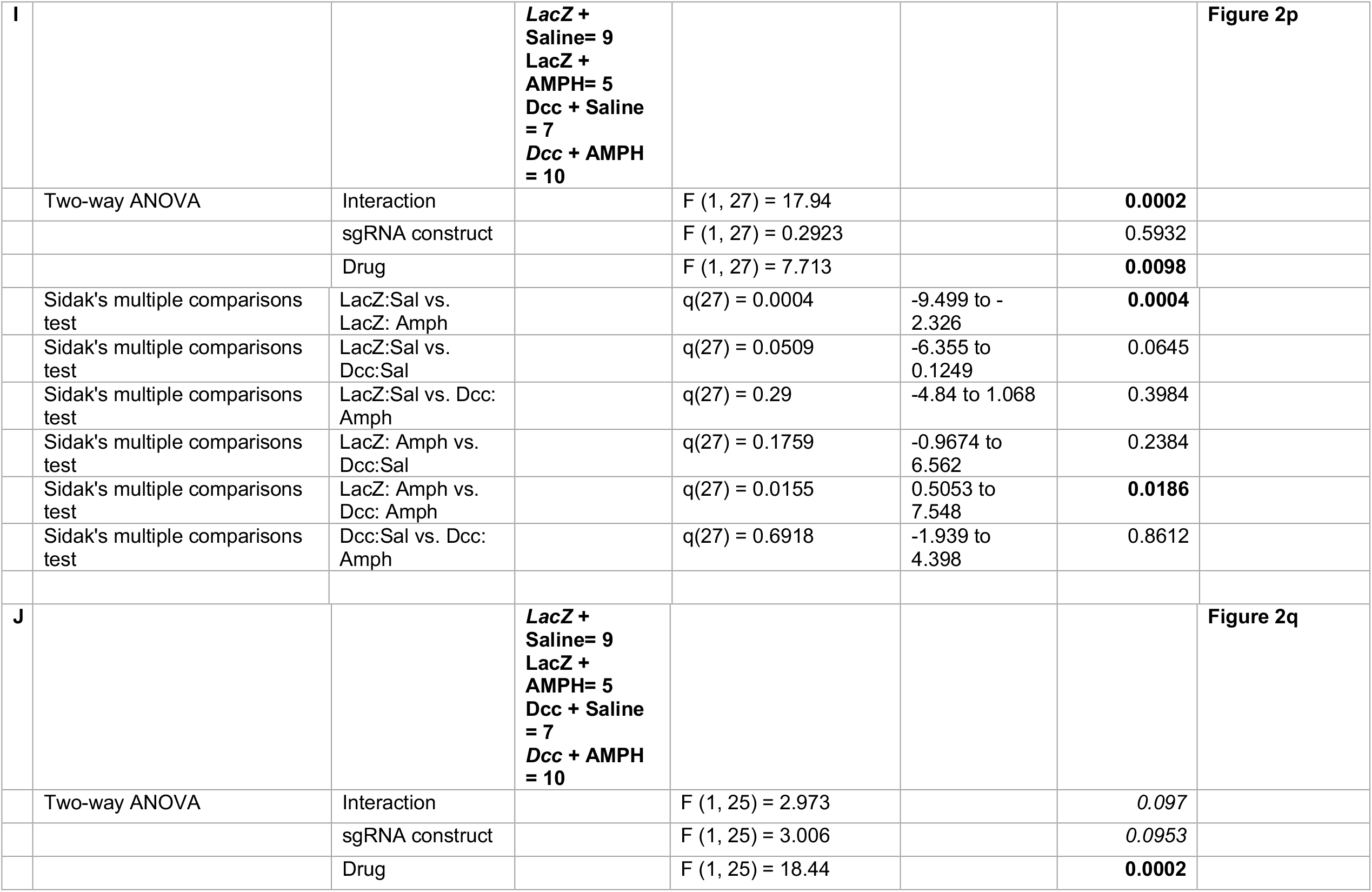
Detailed statistics for Figure 5.

## Discussion

Exposure to a drug is a necessary component to develop addiction or drug-associated psychiatric disorders, but it is not sufficient – only a subset of drug users progress to drug dependence or experience mental and behavioral disorders.^76,77^ What determines why drugs have harmful consequences in some individuals, but not in others, is not well understood. Here we identify for the first time a molecular pathway that is differentially regulated by the same drug experience in adolescent mice, depending on their sex and specific age in adolescence. This different signal encodes the presence or absence of enduring negative outcomes. In early-adolescent males, but not females, a rewarding regimen of 4 mg/kg amphetamine (AMPH) downregulates *Dcc* mRNA expression in dopamine neurons. However, in mid-adolescence, AMPH downregulates *Dcc* mRNA expression in females only. Chronological age and biological sex therefore interact to modulate the impact of drugs of abuse on guidance cue receptors in adolescence. Downregulation of *Dcc* mRNA in dopamine neurons by AMPH in early adolescent males is linked to adult alterations in dopamine innervation to the PFC and to deficits in inhibitory control. Females exposed to AMPH either early or in mid-adolescence do not show these changes. AMPH in mid-adolescent females downregulates *Dcc*, but also leads to a compensatory upregulation of Netrin-1 in the nucleus accumbens, which may actively protect them against deleterious effects. We show that the male-specific deficits in PFC dopamine connectivity and cognitive function in adulthood result from AMPH-induced targeting errors by dopamine axons at the level of the developing NAc, producing incorrect segregation of mesocortical and mesolimbic dopamine projections because mesolimbic dopamine axons end up ectopically innervating the PFC. This effect is absent upon exposure to a therapeutic regimen of amphetamine (Adderall-like dose, ALD) known not to alter *Dcc* mRNA in DA neurons,^62^ reinforcing the idea that *Dcc* downregulation negatively impacts neurodevelopment. Indeed, compensating for the downregulation of *Dcc* via CRISPRa targeted gene therapy, and therefore restoring functional DCC receptor protein levels, prevents adult cognitive impairment in males exposed to AMPH in early adolescence.

Sex differences in addiction risk are well noted, with important disparities between men and women in initiation, escalation, and cessation. For example, adult women are at greater risk than men of quickly progressing to dependence shortly after initiation of cocaine use.^48^ How sex differences in the enduring consequences of drug use are produced in response to the same triggering event remains poorly understood. In this study we found no evidence of a sex effect in the *immediate* rewarding effect of AMPH administration in adolescence since all mice, regardless of age of exposure or sex, show a strong conditioned place preference for AMPH at a dose shown previously to produce peak plasma levels analogous to those seen in recreational users.^62^ In contrast, we found overt sex differences in the *Dcc*-dependent neurodevelopmental impact of AMPH exposure, with females actively protected via compensatory changes in guidance cue expression. However, it is very important not to interpret this result as if females are impervious to any detrimental effects of drugs of abuse in adolescence. While females are protected against *Dcc*-dependent consequences of amphetamine exposure in early- or mid-adolescence, they may still be vulnerable to changes in other physiological systems and behavioral domains, or the effects of other drugs.^78,79^

Inhibitory control is defined as the ability to suppress a given action in response to environmental cues and is known to be sensitive to changes in PFC dopamine tone.^53,65,80^ Deficits in action impulsivity is an endophenotype associated with substance abuse outcomes^56,57,81^ and in adolescents this trait appears to promote the transition from recreational to compulsive drug use.^82,83^ Performance on a Go/No-Go task in youth not only predicts future drug use, but also this association seems to be stronger for teens that are already heavy drug users, suggesting that action impulsivity may predate drug use and be triggered or exacerbated by drug exposure itself.^82^ Action impulsivity is a risk factor for addiction and is also considered a hallmark symptom of attention deficit/hyperactivity disorder (ADHD) which is typically managed with low dose stimulants such as amphetamine (Adderall) or methylphenidate (Ritalin).^84^ Here we show that amphetamine doses equivalent to those used therapeutically in humans do not disrupt adolescent dopamine axon targeting and growth. This is in line with results from previous research in rodents and in non-human primates,^62,71,72^ and with epidemiological evidence showing not only that stimulant medication itself does not increase risk for addiction, but may also counteract the enhanced predisposition for unmedicated ADHD patients to develop substance use disorders later in life. ^83–86^

Our findings provide important mechanistic insight regarding the critical role for the Netrin-1/DCC system in adolescent neurodevelopment,^4,13^ and its strong link to psychiatric disorders of an adolescent onset.^85,86^ Although the regulation of guidance cues by adolescent experience is a nascent field, recent evidence indicates that exposure to AMPH in adolescence is not the only experience that can regulate Netrin-1 and *Dcc* expression. Social defeat stress in adolescence, but not in adulthood, downregulates *Dcc* expression in dopamine neurons in male mice ^35^ and mild traumatic brain injury in adolescent males alters Netrin-1 in the NAc.^87^ Both of these experiential regulations of Netrin-1/*Dcc* expression are associated with alterations in mesocorticolimbic dopamine circuit. Whether positive experiences could alter *Dcc* and/or Netrin-1 levels has yet to be determined but this concept has immense promise for therapeutic applications.

How susceptibility and resilience to addiction is partitioned among drug users remains largely unknown. Understanding how drug use in adolescence produces age- and sex-dependent outcomes is critical to advance addiction research, prevention, and treatment efforts. Here we demonstrate for the first time that exposure to stimulant drugs of abuse in adolescence induce axonal targeting errors, preventing the proper exclusion of mesolimbic dopamine axons from the PFC and leading to cognitive impairments that persist into adulthood. These effects are sex-specific, are mediated by the Netrin-1/DCC guidance cue system, are not observed following therapeutic-like doses, and can be prevented using gene editing strategies. We propose that Netrin-1/DCC signaling functions as a molecular switch to determine whether exposure to the same experience yields to psychiatric vulnerability or resilience.

## Acknowledgements

The authors dedicate this work to the late Dominique Nouel – an excellent and dedicated scientist, a kind mentor and colleague, and a generous and loyal friend. This work was supported by the National Institute on Drug Abuse at the National Institutes of Health (R01DA037911 to C.F.; F31DA041188 to L.M.R.), the Natural Sciences and Engineering Research Council of Canada (RGPIN-2020-04703 to C.F.), the Canadian Institutes of Health Research (FRN: 156272 to C.F.), L.M.R. was also supported by predoctoral fellowships from The Djavad Mowafaghian Foundation and Fulbright Canada, and a post-doctoral NIDA-Inserm Drug Abuse Research Fellowship. The authors declare no conflicts of interest. J.M.R.L was supported by a Doctoral Training fellowship from the Fonds de Recherche du Québec en Santé (JMRL) and a Graduate Student Fellowship from the Healthy Brains for Healthy Lives initiative of the Canada First Research Excellence Fund at McGill University, Canada.

## Author contributions

L.M.R. and C.F. designed the studies. L.M.R., C.P., J.M.R.L., S.I., T.O., and J.G.E. performed behavioral experiments. G.H., D.N., S.C., J.M.R-L, and M.G. performed qPCR and western blotting experiments. L.M.R., D.M., D.N., J.Z., and M-Y. H. performed stereological experiments. M.P., M.W. and B.K. performed Golgi experiments. G.H. and S.B.N. performed CRISPR validation studies, with in vitro work done in the lab of L-É. T. J.J.D. provided CRISPRa virus material and the sgRNAs, based on sgRNAs designed by K.S. and L.M.R. G.H. performed CRISPR virus injections and behavioral experiments. L.M.R, G.H., C.P., and C.F. analysed the data. L.M.R. and C.F. wrote the manuscript. All authors discussed the results, edited, and approved the manuscript.

## Notes

### Competing Interest Statement

The authors have declared no competing interest.

## References

1. Sawyer, S. M., Azzopardi, P. S., Wickremarathne, D. & Patton, G. C. The age of adolescence. Lancet Child Adolesc Heal 2, 223–228 (2018).

2. Schneider, M. Adolescence as a vulnerable period to alter rodent behavior. Cell Tissue Res 354, (2013).

3. Granata, L., Gildawie, K. R., Ismail, N., Brenhouse, H. C. & Kopec, A. M. Immune signaling as a node of interaction between systems that sex-specifically develop during puberty and adolescence. Dev Cogn Neuros-neth 57, 101143 (2022).

4. Reynolds, L. M. & Flores, C. Mesocorticolimbic Dopamine Pathways Across Adolescence: Diversity in Development. Front Neural Circuit 15, 735625 (2021).

5. Spear, L. P. The adolescent brain and age-related behavioral manifestations. Neurosci Biobehav Rev 24, 417 463 (2000).

6. Larsen, B. et al. Maturation of the human striatal dopamine system revealed by PET and quantitative MRI. Nat Commun 11, 846 (2020).

7. Wahlstrom, D., White, T. & Luciana, M. Neurobehavioral evidence for changes in dopamine system activity during adolescence. Neurosci Biobehav Rev 34, 631 648 (2010).

8. Luciana, M., Wahlstrom, D., Porter, J. N. & Collins, P. F. Dopaminergic modulation of incentive motivation in adolescence: age-related changes in signaling, individual differences, and implications for the development of self-regulation. Dev Psychol 48, 844 861 (2012).

9. Rothmond, D. A., Weickert, C. S. & Webster, M. J. Developmental changes in human dopamine neurotransmission: cortical receptors and terminators. Bmc Neurosci 13, 18 (2012).

10. Weickert, C. S. et al. Postnatal alterations in dopaminergic markers in the human prefrontal cortex. Neuroscience 144, 1109 1119 (2007).

11. Rosenberg, D. R. & Lewis, D. A. Changes in the dopaminergic innervation of monkey prefrontal cortex during late postnatal development: a tyrosine hydroxylase immunohistochemical study. Biol Psychiat 36, 272 277 (1994).

12. Rosenberg, D. R. & Lewis, D. A. Postnatal maturation of the dopaminergic innervation of monkey prefrontal and motor cortices: a tyrosine hydroxylase immunohistochemical analysis. J Comp Neurol 358, 383 400 (1995).

13. Reynolds, L. M. & Flores, C. Adolescent dopamine development: connecting experience with vulnerability or resilience to psychiatric disease. in Diagnosis, Management and Modeling of Neurodevelopmental Disorders (eds. Martin, C. R., Preedy, V. R. & Rajendram, R.) (Academic Press, 2021). doi:10.1016/b978-0-12-817988-8.00026-9.

14. Hoops, D. & Flores, C. Making Dopamine Connections in Adolescence. Trends Neurosci 40, (2017).

15. Barth, B., Portella, A. K., Dubé, L., Meaney, M. J. & Silveira, P. P. Early Life Origins of Ageing and Longevity. in Early Life Origins of Ageing and Longevity 121–140 (2019). doi:10.1007/978-3-030-24958-8_7.

16. Sebastian, C., Viding, E., Williams, K. D. & Blakemore, S.-J. Social brain development and the affective consequences of ostracism in adolescence. Brain Cognition 72, 134–145 (2010).

17. Orben, A., Tomova, L. & Blakemore, S.-J. The effects of social deprivation on adolescent development and mental health. Lancet Child Adolesc Heal 4, 634–640 (2020).

18. Blakemore, S.-J. The social brain in adolescence. Nat Rev Neurosci 9, 267–277 (2008).

19. Baarendse, P. J. J., Counotte, D. S., O’Donnell, P. & Vanderschuren, L. J. M. J. Early social experience is critical for the development of cognitive control and dopamine modulation of prefrontal cortex function. Neuropsychopharmacol 38, 1485 1494 (2013).

20. Yamamuro, K. et al. Social Isolation During the Critical Period Reduces Synaptic and Intrinsic Excitability of a Subtype of Pyramidal Cell in Mouse Prefrontal Cortex. Cereb Cortex 28, 1 13 (2017).

21. Makinodan, M., Rosen, K. M., Ito, S. & Corfas, G. A critical period for social experience-dependent oligodendrocyte maturation and myelination. Science 337, 1357 1360 (2012).

22. Berding, K. et al. Diet and the Microbiota–Gut–Brain Axis: Sowing the Seeds of Good Mental Health. Adv Nutr 12, nmaa181- (2021).

23. Neufeld, K.-A. M., Luczynski, P., Oriach, C. S., Dinan, T. G. & Cryan, J. F. What’s bugging your teen?—The microbiota and adolescent mental health. Neurosci Biobehav Rev 70, 300–312 (2016).

24. Graupensperger, S., Sutcliffe, J. & Vella, S. A. Prospective Associations between Sport Participation and Indices of Mental Health across Adolescence. J Youth Adolescence 50, 1450–1463 (2021).

25. Boone, E. M. & Leadbeater, B. J. Game On: Diminishing Risks for Depressive Symptoms in Early Adolescence Through Positive Involvement in Team Sports. J Res Adolescence 16, 79–90 (2006).

26. Hopkins, M. E., Nitecki, R. & Bucci, D. J. Physical exercise during adolescence versus adulthood: differential effects on object recognition memory and brain-derived neurotrophic factor levels. Neuroscience 194, 84–94 (2011).

27. Gama, C. S. et al. Effects of omega-3 dietary supplement in prevention of positive, negative and cognitive symptoms: a study in adolescent rats with ketamine-induced model of schizophrenia. Schizophr Res 141, 162 167 (2012).

28. Serafine, K. M., Labay, C. & France, C. P. Dietary supplementation with fish oil prevents high fat diet-induced enhancement of sensitivity to the locomotor stimulating effects of cocaine in adolescent female rats. Drug Alcohol Depen 165, 45 52 (2016).

29. Eddy, M. C. & Green, J. T. Running Wheel Exercise Reduces Renewal of Extinguished Instrumental Behavior and Alters Medial Prefrontal Cortex Neurons in Adolescent, But Not Adult, Rats. Behav Neurosci 131, 460–469 (2017).

30. Rijlaarsdam, J., Cecil, C. A. M., Buil, J. M., Lier, P. A. C. van & Barker, E. D. Exposure to Bullying and General Psychopathology: A Prospective, Longitudinal Study. Res Child Adolesc Psychopathol 49, 727–736 (2021).

31. Compas, B. E., Orosan, P. G. & Grant, K. E. Adolescent stress and coping: implications for psychopathology during adolescence. J Adolescence 16, 331–349 (1993).

32. Sheth, C., McGlade, E. & Yurgelun-Todd, D. Chronic Stress in Adolescents and Its Neurobiological and Psychopathological Consequences: An RDoC Perspective. Chronic Stress 1, 2470547017715645 (2017).

33. Novick, A. M., Forster, G. L., Tejani-Butt, S. M. & Watt, M. J. Adolescent social defeat alters markers of adult dopaminergic function. Brain Res Bull 86, 123 128 (2011).

34. Watt, M. J. et al. Decreased prefrontal cortex dopamine activity following adolescent social defeat in male rats: role of dopamine D2 receptors. Psychopharmacology 231, (2013).

35. Vassilev, P. et al. Unique effects of social defeat stress in adolescent male mice on the Netrin-1/DCC pathway, prefrontal cortex dopamine and cognition (Social stress in adolescent vs. adult male mice). Eneuro ENEURO.0045-21.2021 (2021) doi:10.1523/eneuro.0045-21.2021.

36. Jordan, C. J. & Andersen, S. L. Sensitive periods of substance abuse: Early risk for the transition to dependence. Dev Cogn Neuros-neth 25, 29 44 (2017).

37. Gulley, J. M. & Juraska, J. M. The effects of abused drugs on adolescent development of corticolimbic circuitry and behavior. Neuroscience 249, (2013).

38. Grant, B. F. Age at smoking onset and its association with alcohol consumption and DSM-IV alcohol abuse and dependence: Results from the national longitudinal alcohol epidemiologic survey. J Subst Abuse 10, 59–73 (1998).

39. Grant, B. F. & Dawson, D. A. Age of onset of drug use and its association with DSM-IV drug abuse and dependence: Results from the national longitudinal alcohol epidemiologic survey. J Subst Abuse 10, 163–173 (1998).

40. Robins, L. N. & Przybeck, T. R. Age of onset of drug use as a factor in drug and other disorders. NIDA research monograph 56, 178 192 (1985).

41. Anthony, J. C. & Petronis, K. R. Early-onset drug use and risk of later drug problems. Drug Alcohol Depen 40, 9–15 (1995).

42. McCabe, S. E., West, B. T., Morales, M., Cranford, J. A. & Boyd, C. J. Does early onset of non-medical use of prescription drugs predict subsequent prescription drug abuse and dependence? Results from a national study. Addiction 102, 1920 1930 (2007).

43. Vega, W. A. et al. Prevalence and age of onset for drug use in seven international sites: results from the international consortium of psychiatric epidemiology. Drug Alcohol Depen 68, 285–297 (2002).

44. Cotto, J. H. et al. Gender effects on drug use, abuse, and dependence: A special analysis of results from the national survey on drug use and health. Gender Med 7, 402–413 (2010).

45. Shansky, R. M. & Murphy, A. Z. Considering sex as a biological variable will require a global shift in science culture. Nat Neurosci 1–8 (2021) doi:10.1038/s41593-021-00806-8.

46. McHugh, R. K., Votaw, V. R., Sugarman, D. E. & Greenfield, S. F. Sex and gender differences in substance use disorders. Clin Psychol Rev 66, (2017).

47. Hernandez-Avila, C. A., Rounsaville, B. J. & Kranzler, H. R. Opioid-, cannabis- and alcohol-dependent women show more rapid progression to substance abuse treatment. Drug Alcohol Depen 74, 265–272 (2004).

48. O’Brien, M. S. & Anthony, J. C. Risk of becoming cocaine dependent: epidemiological estimates for the United States, 2000-2001. Neuropsychopharmacol 30, 1006 1018 (2005).

49. Becker, J. B. & Chartoff, E. Sex differences in neural mechanisms mediating reward and addiction. Neuropsychopharmacol 26, 1 (2018).

50. Moore, S. W., Tessier-Lavigne, M. & Kennedy, T. E. Netrins and their receptors. Adv Exp Med Biol 621, 17 31 (2007).

51. Boyer, N. P. & Gupton, S. L. Revisiting Netrin-1: One Who Guides (Axons). Front Cell Neurosci 12, 221 (2018).

52. Bradford, D., Cole, S. J. & Cooper, H. M. Netrin-1: diversity in development. Int J Biochem Cell Biology 41, 487 493 (2009).

53. Reynolds, L. M. et al. DCC Receptors Drive Prefrontal Cortex Maturation by Determining Dopamine Axon Targeting in Adolescence. Biol Psychiat 83, 181 192 (2018).

54. Hoops, D., Reynolds, L. M., Restrepo-Lozano, J. M. & Flores, C. Dopamine Development in the Mouse Orbital Prefrontal Cortex Is Protracted and Sensitive to Amphetamine in Adolescence. Eneuro 5, ENEURO.0372-17.2017 (2018).

55. Manitt, C. et al. dcc orchestrates the development of the prefrontal cortex during adolescence and is altered in psychiatric patients. Transl Psychiat 3, e338 (2013).

56. Jentsch, J. D. et al. Dissecting impulsivity and its relationships to drug addictions. Ann Ny Acad Sci 1327, 1 26 (2014).

57. Bari, A. & Robbins, T. W. Inhibition and impulsivity: behavioral and neural basis of response control. Prog Neurobiol 108, 44 79 (2013).

58. Cuesta, S. et al. Non-Contingent Exposure to Amphetamine in Adolescence Recruits miR-218 to Regulate Dcc Expression in the VTA. Neuropsychopharmacol 43, 900 911 (2018).

59. Phillips, R. A. et al. An atlas of transcriptionally defined cell populations in the rat ventral tegmental area. Cell Reports 39, 110616 (2022).

60. Manitt, C. et al. Peri-pubertal emergence of UNC-5 homologue expression by dopamine neurons in rodents. Plos One 5, e11463 (2010).

61. Cuesta, S. et al. Dopamine Axon Targeting in the Nucleus Accumbens in Adolescence Requires Netrin-1. Frontiers Cell Dev Biology 8, 487 (2020).

62. Cuesta, S. et al. DCC-related developmental effects of abused-versus therapeutic-like amphetamine doses in adolescence. Addict Biol e12791 (2019) doi:10.1111/adb.12791.

63. Yetnikoff, L., Pokinko, M., Arvanitogiannis, A. & Flores, C. Adolescence: a time of transition for the phenotype of dcc heterozygous mice. Psychopharmacology 231, 1705 1714 (2014).

64. Reynolds, L. M. et al. Amphetamine in Adolescence Disrupts the Development of Medial Prefrontal Cortex Dopamine Connectivity in a dcc-Dependent Manner. Neuropsychopharmacol 40, 1101 1112 (2015).

65. Reynolds, L. M. et al. Early Adolescence is a Critical Period for the Maturation of Inhibitory Behavior. Cereb Cortex 29, 3676–3686 (2019).

66. Manitt, C. et al. The netrin receptor DCC is required in the pubertal organization of mesocortical dopamine circuitry. J Neurosci 31, 8381 8394 (2011).

67. Manitt, C. & Kennedy, T. E. Chapter 32 Where the rubber meets the road: netrin expression and function in developing and adult nervous systems. Prog Brain Res 137, 425–442 (2002).

68. Varadarajan, S. G. et al. Netrin1 Produced by Neural Progenitors, Not Floor Plate Cells, Is Required for Axon Guidance in the Spinal Cord. Neuron 94, (2017).

69. Wu, Z. et al. Long-Range Guidance of Spinal Commissural Axons by Netrin1 and Sonic Hedgehog from Midline Floor Plate Cells. Neuron 101, 635-647.e4 (2019).

70. Moreno-Bravo, J. A., Puiggros, S. R., Mehlen, P. & Chédotal, A. Synergistic Activity of Floor-Plate- and Ventricular-Zone-Derived Netrin-1 in Spinal Cord Commissural Axon Guidance. Neuron 101, 625-634.e3 (2019).

71. Labonté, B. et al. Adolescent amphetamine exposure elicits dose-specific effects on monoaminergic neurotransmission and behaviour in adulthood. Int J Neuropsychoph 15, 1319 1330 (2012).

72. Soto, P. L. et al. Long-Term Exposure to Oral Methylphenidate or dl-Amphetamine Mixture in Peri-Adolescent Rhesus Monkeys: Effects on Physiology, Behavior, and Dopamine System Development. Neuropsychopharmacol 37, 2566 2579 (2012).

73. Cho, K. R. et al. The DCC gene: structural analysis and mutations in colorectal carcinomas. Genomics 19, 525 531 (1994).

74. Reale, M. A. et al. Expression and alternative splicing of the deleted in colorectal cancer (DCC) gene in normal and malignant tissues. Cancer Res 54, 4493–501 (1994).

75. Savell, K. E. et al. A Neuron-Optimized CRISPR/dCas9 Activation System for Robust and Specific Gene Regulation. Eneuro 6, ENEURO.0495-18.2019 (2019).

76. Anthony, J. C., Warner, L. A. & Kessler, R. C. Comparative Epidemiology of Dependence on Tobacco, Alcohol, Controlled Substances, and Inhalants: Basic Findings From the National Comorbidity Survey. Exp Clin Psychopharm 2, 244–268 (1994).

77. Pedersen, C. B. et al. A Comprehensive Nationwide Study of the Incidence Rate and Lifetime Risk for Treated Mental Disorders. Jama Psychiat 71, 573–581 (2014).

78. Prini, P. et al. Adolescent THC exposure in female rats leads to cognitive deficits through a mechanism involving chromatin modifications in the prefrontal cortex. J Psychiatr Neurosci 43, 87–101 (2017).

79. Rubino, T. et al. Adolescent exposure to THC in female rats disrupts developmental changes in the prefrontal cortex. Neurobiol Dis 73, 60 69 (2015).

80. Yan, R., Wang, T. & Zhou, Q. Elevated dopamine signaling from ventral tegmental area to prefrontal cortical parvalbumin neurons drives conditioned inhibition. Proc National Acad Sci 116, 13077–13086 (2019).

81. Grant, J. E. & Chamberlain, S. R. Impulsive action and impulsive choice across substance and behavioral addictions: Cause or consequence? Addict Behav 39, 1632–1639 (2014).

82. Mahmood, O. M. et al. Adolescents’ fMRI activation to a response inhibition task predicts future substance use. Addict Behav 38, 1435 1441 (2013).

83. Nigg, J. T. et al. Poor Response Inhibition as a Predictor of Problem Drinking and Illicit Drug Use in Adolescents at Risk for Alcoholism and Other Substance Use Disorders. J Am Acad Child Adolesc Psychiatry 45, 468–475 (2006).

84. Groman, S. M., James, A. S. & Jentsch, J. D. Poor response inhibition: At the nexus between substance abuse and attention deficit/hyperactivity disorder. Neurosci Biobehav Rev 33, 690–698 (2009).

85. Vosberg, D. E., Leyton, M. & Flores, C. The Netrin-1/DCC guidance system: dopamine pathway maturation and psychiatric disorders emerging in adolescence. Mol Psychiatr 1–11 (2019) doi:10.1038/s41380-019-0561-7.

86. Torres-Berrío, A., Hernandez, G., Nestler, E. J. & Flores, C. The Netrin-1/DCC guidance cue pathway as a molecular target in depression: Translational evidence. Biol Psychiat (2020) doi:10.1016/j.biopsych.2020.04.025.

87. Kaukas, L., Holmes, J. L., Rahimi, F., Collins-Praino, L. & Corrigan, F. Injury during adolescence leads to sex-specific executive function deficits in adulthood in a pre-clinical model of mild traumatic brain injury. Behav Brain Res 113067 (2020) doi:10.1016/j.bbr.2020.113067.

88. Yetnikoff, L., Almey, A., Arvanitogiannis, A. & Flores, C. Abolition of the behavioral phenotype of adult netrin-1 receptor deficient mice by exposure to amphetamine during the juvenile period. Psychopharmacology 217, 505 514 (2011).

89. Keino-Masu, K. et al. Deleted in Colorectal Cancer (DCC) Encodes a Netrin Receptor. Cell 87, 175–185 (1996).

90. Cooper, H. M., Armes, P., Britto, J., Gad, J. & Wilks, A. F. Cloning of the mouse homologue of the deleted in colorectal cancer gene (mDCC) and its expression in the developing mouse embryo. Oncogene 11, 2243–54 (1995).

