## Supplementary data and methods for "Amphetamine disrupts dopamine axon growth in adolescence by a sex-specific mechanism"

Supplementary Information

### Materials and Methods

Animals

Experimental procedures were performed according to the guidelines of the Canadian Council of Animal Care and approved by the McGill University/Douglas Mental Health University Institute Animal Care Committee. $DAT^{Cre}$ or wildtype C57BL/6J mice were bred in the Douglas Mental Health University Institute Neurophenotyping center, or were obtained from Charles River Laboratories (Saint-Constant, QC, Canada). All mice were maintained on a 12-h light–dark cycle (light on at 0800 h) and given *ad libitum* access to food and water unless otherwise stated. Male and female mice were housed with same-sex littermates throughout the experimental procedures.

Drugs and dose

d-Amphetamine sulfate (AMPH; Sigma-Aldrich, Dorset, United Kingdom) was dissolved in 0.9% saline. All AMPH injections were administered i.p. at a volume of 0.1ml/10g. A 'recreational-like' dose of 4 mg/kg was used to achieve peak plasma AMPH levels of 1300 ± 79 ng/mL 5 minutes post-injection, consistent with plasma levels induced by recreational use of AMPH in humans. A low, 'Adderall-like' dose (ALD) of 0.5 mg/kg was used to achieve peak plasma levels of 97 ± 21 ng/mL, in line with those observed following therapeutic administration in humans.[1]

*AMPH and ALD treatment regimen*. Mice received one injection of AMPH or ALD (experimental group) or saline (control group), once every other day for a total of 5 treatment days. This treatment regimen was administered either during early adolescence (from PND 22±1 to PND 31±1), or during mid-adolescence (PND35±1 to PND 44±1). Locomotor activity was measured 15min prior to and 90min after each AMPH, ALD, or saline injection.

Axon-initiated recombination

We tracked the growth of dopamine axons across adolescence using axon-initiated recombination.[2] Importantly, we used $DAT^{Cre}$ mice and modified the viruses used in these experiments in order to produce cell-type specific labeling confined only to dopamine neurons. At PND21, we injected a retrogradely transported virus expressing a Cre-dependent Flp recombinase (CAV-FLEX-Flp, BioCampus Montpellier) unilaterally into the NAcc of $DAT^{Cre}$ mice. This design limits expression of the Flp recombinase to DAT-expressing (i.e. dopaminergic) neurons that project to the NAcc at PND 21. Simultaneously, we injected a Flp-dependent enhanced yellow fluorescent protein (eYFP) virus fDIO-eYFP (pAAV-Ef1a-fDIO-EYFP-WPRE-pA, UNC Vector Core) into the ipsilateral VTA. Thus, eYFP will only be expressed in a projection-specific and cell-type specific manner.

Stereology
All procedures have been previously reported in detail.[3] Briefly, mice were perfused at PND75±15 with 4% paraformaldehyde, and their brains sliced into 35μm sections using a Leica vibratome. Sections were incubated for 48 hours with a polyclonal anti-GFP raised in chicken (1:1000, antibody #1020, Aves labs) and a polyclonal rabbit anti-tyrosine hydroxylase (TH) antibody (1:1000, AB152, Millipore Bioscience Research Reagents). Immunostaining was visualized with Alexa Fluor 488- and Alexa Fluor 594-conjugated secondary antibodies raised in goat (1:500; Invitrogen).

We used Stereoinvestigator (MBF, St. Albans VT) to quantify (a) the span and density of TH-positive innervation, (b) the number of TH-positive, eYFP-positive varicosities, and (c) the number of TH-negative, eYFP-positive varicosities in the nucleus accumbens (NAc) and the cingulate (Cg1), prelimbic (PrL), and infralimbic (IL) subregions of the pregenual prefrontal cortex (PFC). We also quantify (a) the number of TH-positive neurons, (b) The number of eYFP-labeled TH positive neurons, and (c) the number of TH-negative, eYFP-positive neurons in the ventral tegmental area (VTA) and substantia nigra pars compacta (SNc).

Innervation volume in cubic micrometers was assessed with the Cavalieri method in Stereoinvestigator. Cells and terminals were quantified using the optical fractionator probe. As in all our previous neuroanatomical studies, we obtained counts only from the right hemisphere because of the lateralization of dopamine systems and the unilateral injections for axon tracing experiments. The Coefficient of error for TH-positive varicosities/cells was below 0.10 for all regions of interest. Counts were performed blind.

*PFC:* Cg1, PrL, and IL subregions of the PFC were delineated according to plates spanning 14 – 18 of the mouse brain atlas[4] and contours were traced at 5X magnification using a Leica DM400B microscope along the dense TH-positive innervation of PFC layers V-VI. An unbiased counting frame (50 x 50 μm) was superimposed on each contour and counts were made at regular predetermined intervals (x= 175 μm, y= 175 μm) from a random start point. TH positive and eYFP positive varicosities were counted at 100X magnification on 5 sections contained within the rostrocaudal borders of our region of interest (Plates 14-18; 1:4 series). A guard zone of 4 μm was used and the optical disector height was set to 10 μm.

*NAc:* An unbiased counting frame (10 x 10 μm) was superimposed on the contour of the NAc and counts were made at regular predetermined intervals (x= 400 μm, y= 400 μm) from a random start point. Counting was performed at 100X magnification on four of the eight sections contained within the rostrocaudal borders of our region of interest (Plates 15–18, 1:4 series). A guard zone of 4μm was used and the optical disector height was set to 5μm.

*Midbrain analysis*: The counting scheme used a 60 × 60 µm counting frame (x =150 µm, y = 150 µm intervals) with a random start point. Counting was performed at 40X magnification in a 1:4 series. A 3 µm guard zone and a probe depth of 10 µm were used. Stereological counts of eYFP and TH co-labeled neuron populations were expressed as proportions.

CRISPR activation

*CRISPR/dCas9 and sgRNA construct design:*
Single guide RNAs compatible with CRISPR activation (CRISPRa) were designed as previously described.[5] Briefly, sgRNA targets were designed using online tools provided by the Zhang Lab at MIT (crispr.mit.edu) and CHOPCHOP (RRID:SCR_015723; http://chopchop.cbu.uib.no/)[6,7] to target within –1000/ -500 bp of the transcription start site (TSS) of the mouse *Dcc* gene. To ensure specificity, all CRISPR RNA (crRNA) sequences were then analyzed with National Center for Biotechnology Information's (NCBI) Basic Local Alignment Search Tool (BLAST). A list of the target sequences is provided in Extended Data Table 1. Custom crRNAs were ordered as oligonucleotide sequences (Sigma Aldrich) with 5' 4-bp overhangs (CACC for the sense strand, AAAC for the antisense strand). crRNAs were annealed, phosphorylated with PNK (NEB), and ligated using T4 ligase (NEB) into the short guide RNA (sgRNA) scaffold using the BbsI cut sites with unique overhangs mentioned above. For crRNA sequences that did not begin with a guanine, the first base of the crRNA sequence was substituted to guanine to maintain compatibility with the U6 promoter. CRISPRa experiments used lentivirus compatible plasmid constructs previously optimized for robust neuronal expression (lenti SYN-FLAG-dCas9-VPR, RRID:Addgene_114196; lenti U6-sgRNA/EF1a-mCherry, RRID:Addgene_114199) [5]. The bacterial LacZ gene target was used as a sgRNA non-targeting control.[8]

*Lentivirus preparation:*
Plasmid preparation. One Shot Stbl3 Chemically Competent E. coli (Invitrogen, Catalog number: C737303), were heat shock transformed to amplify all plasmids. Plasmids were purified using a Qiagen EndoFree Plasmid Maxi Kit (Catalog number: 12362).
Lentivirus production . Viruses were produced in a sterile environment subject to BSL-2 safety by transfecting HEK293T cells with specified CRISPR-dCas9 plasmids, the psPAX2 packaging plasmid, and the pCMV-VSV G envelope plasmid (Addgene plasmids #12260 and #8454) with FuGene HD (Promega) for 48 hours. Cells were incubated at 37°C and 5% $CO_2$ in supplemented Ultraculture media (L-glutamine, sodium pyruvate, and sodium bicarbonate) in either a T75 or T225 culture flask. Viruses were purified from the supernatant using filter (0.45mm) and ultracentrifugation (25,000 rpm, 1 h 45 min at 4°C) . Viral titer was determined using a qPCR Lentivirus Titration kit (Lenti-X, qRT-PCR Titration kit, Takara). After 40–48 h, lentiviruses were concentrated with Lenti-X concentrator (Takara), resuspended in sterile PBS, and used or frozen at -80°C immediately. Only viruses with a titer of > 1 x 10^15 GC/ml were used. Viruses were stored in sterile PBS at 80°C in single-use aliquots.

*In vitro validation:*
Primary mesencephalic neuron cultures were prepared from dissections of male and female postnatal day 0-2 (P0 to P2) C57/BL6J mice according to a protocol described previously.[9] Briefly, mice were cryoanesthetized and the brain was rapidly obtained to isolate the VTA and SNc. The tissue was digested with papain and triturated to obtain a single-cell suspension. The cells were plated on 15 mm diameter glass coverslips at 120 000 cells/ml on top of a pre-established cortical astrocyte layer.

For immunofluorescent imaging, neuronal cultures were fixed with 4% paraformaldehyde (PFA), permeabilized, and nonspecific binding sites were blocked using BSA. Dopamine neurons were identified by immunofluorescence using a primary anti-TH antibody (1:1000, AB152, Millipore Sigma, USA). Cultures were washed with PBS and incubated for 2h at room temperature with a secondary antibody (anti-mouse Alexa Fluor-488, 1:1000, Invitrogen) (green in panel 1E). Expression of the virally expressed sgRNAs and dCAS9 was validated by detecting the associated co-expressed mCherry protein (purple in panel 1E). Finally, coverslips were washed, counterstained with DAPI (blue) and mounted in Fluoromount-G (Southern Biotech) on Superfrost/Plus microscope slides. For qPCR experiments, mRNA was extracted with trizol from the cells on the cover slips.

*In vivo validation:*
Early adolescent male mice were bilaterally infused with 1.0 µl of total lentivirus mix with 0.33 µl of the 4 sgRNAs and 0.66 µl of the dCas9-VPR virus in sterile PBS.[5] Viral transduction and *Dcc* mRNA overexpression were assessed 10 days later via immunofluorescence and qPCR in the VTA. DCC protein expression was assessed by western blot analysis in the NAc, where DCC protein is only expressed in dopamine axons.[10]

*Experimental design:*
Male mice received VTA stereotaxic bilateral infusions of the dCas9-VPR and *Dcc* targeting sgRNAs or (control) dCas9-VPR and *LacZ* sgRNA lentiviral constructs at P21. Two days later, they began the AMPH or saline treatment regimen. In adulthood, mice were tested in the Go/No-Go task.

*Infection and probe placement verification:*
Adult mice received an overdose of ketamine (50 mg/kg), xylazine (5 mg/kg), and acepromazine (1 mg/kg) through intraperitoneal injection and were perfused intracardially with ice-cold phosphate-buffered saline (PBS, 1x) followed by ice-cold 4% paraformaldehyde (PFA, pH = 7.4). Brains were dissected, post-fixed in 4% PFA overnight at 4°C, and transferred to 1x PBS 24 hours before slicing. 35µm coronal sections were obtained using a vibratome (Leica Biosystems VT1000S) and stored in a cryo-protective solution at -20°C until processing.

Every second section was processed for visualization of TH+ neurons and mCherry. Sections were rinsed three times for 10 minutes with 1x PBS and blocked in 2% bovine serum albumin (in 1x PBS and Tween-20) for 1 hour at room temperature. Sections were then incubated in primary antibodies, including mouse anti-TH (Millipore Sigma, cat. no. MAB318) and rabbit anti-RFP (Rockland, cat. no. 600-401-379), for 48 hours at 4°C. Sections were rinsed three times for 10 min with 1x PBS and incubated in secondary antibodies, including goat anti-mouse Alexa Fluor 488 (Invitrogen, cat. no. A-11001) and donkey anti-rabbit Alexa Fluor 594 (Invitrogen, cat. no. A-21207), for 1 hour at room temperature. Sections were rinsed three times in 1x PBS and mounted with VECTASHIELD Hardset antifade mounting medium with DAPI (Vector Laboratories, cat. no. H-1500-10). Representative images were taken using the Stereo Investigator software (MBF Bioscience) with an epifluorescent microscope (Leica DM400X3).

<u>Morphological analysis of PFC pyramidal neurons</u>

*Golgi–Cox staining*. Mice were deeply anesthetized with sodium pentobarbital (>75 mg/kg; i.p.) and perfused with 0.9% saline, their brains were then processed for Golgi-Cox staining as previously.[2,10,11]

*Anatomical analysis.* Basilar dendritic arbors and spines of layer V mPFC pyramidal neurons were analyzed to quantify the total arbor length, number of branches, and spine density of each cell. Neurons from the Cg1, PrL, and IL subregions of the pregenual PFC were analyzed. A Leica model DM400 microscope equipped with a Ludl XYZ motorized stage was used to identify cells, trace dendritic arbors, and quantify dendritic spines. Relevant regions were first identified at low magnification (5X objective). Cells that were chosen for tracing and analysis were required to have intact branches, well impregnated staining, and not obscured by blood vessels, astrocytes, or heavy clusters of dendrites from other cells. Neurolucida software (MicroBrightField) was used to trace the dendritic arbors of selected cells and to quantify dendritic arbor length, dendrite number, and the spine density on selected dendrite segments. For both dendritic arbor and spine density analysis, the same neurons were sampled. One dendritic segment (third-order tip or greater) was analyzed per neuron under the 100X objective. Spines were always counted from the last branch point to the terminal tip of the dendrite. No attempt was made to correct for the fact that some spines are obscured from view, so the measure of spine density necessarily underestimates total spine density. Anatomical analysis was conducted blind to treatment condition. A minimum of four cells were analyzed per brain, and averaged across each subject.

<u>Behavior</u>

*Go/No-Go.* The Go/No-Go task was performed as previously.[2] Briefly, food-restricted mice (85% free feeding weight) were trained to nosepoke for chocolate-flavored dustless precision pellets (BioServ, Inc., Flemington, NJ, USA) in MedAssociates operant boxes. The mice first undergo discrimination training, where they learn to nose poke only when signaled to do so. Premature responses in discrimination training sessions are a measure of waiting impulsivity. After training, mice underwent daily sessions of the Go/No-Go Task, which requires the mice to respond to a lighted 'Go' cue or inhibit their response to this cue when presented in tandem with an auditory 'No-Go' cue. Within each session, the number of 'Go' and 'No-Go' trials were given in an approximately 1:1 ratio and presented in a randomized order. Each session lasted 30 minutes and consisted of approximately 30-50 'Go' and 30-50 'No-Go' trials. Number of responses to the No-Go cue (commission errors) and correct responses to the Go cue (hits) were analyzed. Commission errors represent a measure of action impulsivity, defined as a failure to appropriately inhibit behavior.

*Conditioned Place Preference.* Male or female mice were tested for conditioned place preference to 4 mg/kg AMPH in early or mid- adolescence. On day 1 mice were allowed to freely explore the CPP apparatus for 30 min, which consisted of 2 distinct chambers (one striped, one polka-dotted) and a neutral (grey) area connecting the chambers. Time spent in each chamber was measured to determine a preference percentage between the chambers for each individual animal, and a biased design was used, i.e. the less preferred chamber during the pretest would be paired with AMPH (experimental group), or with saline (control group). Following the pretest day, animals were exclusively exposed to one chamber, paired either with an AMPH (experimental group) or saline (control group) injection, for 30 minutes every other day for 9 treatment days. This is identical to the treatment regimen used in all anatomical, behavioral, and neurochemical experiments in this study. After the last day of injections, mice were once again allowed to freely explore the full enclosure for a 20 minute post-test while the time spent in each was measured. A delta preference score was then calculated for each mouse by subtracting the time spent in the originally unpreferred chamber during pretest from the time spent in that same chamber during post-test, *Place Preference = time POST - time PRE*.

Quantitative Real-Time PCR

qPCR experiments were performed as previously described.[12] Briefly, mice were euthanized one week following the end of amphetamine treatment, their brains removed and rapidly frozen in dry ice-cooled 2-methylbutane (Fisher Scientific, Hampton, NH, USA). Brains were sliced in 1-mm-thick coronal slices using a cryostat and VTA, NAcc, and mPFC punches were taken from the resulting sections. Total RNA and microRNA were extracted from the VTA punches using an mRNAeasy Micro Kit (Qiagen). *Dcc* mRNA was reverse transcribed using a High-Capacity cDNA Reverse Transcription Kit (Applied Biosystems), and real-time PCR was performed using a TaqMan assay kit (Applied Biosystems) on a 7900HT RT PCR system (Applied Biosystems) in technical triplicates. *Gapdh* was used as a reference gene to control for experimental variability. A TaqMan MicroRNA Reverse Transcription Kit was used alongside the corresponding miRNA TaqMan probes (Applied Biosystems, Foster City, CA) to reverse transcribe and perform Real-Time PCR for miR-218, and expression levels were calculated using the AQ standard curve method. The small nucleolar RNA (snoRNA) RNU6B was used as an endogenous control to normalize miR-218 expression.

Western blot

Whole brains from PND21 and 35 male and female C57BL/6J mice were flash frozen in 2-methylbutane. Bilateral punches from the NAc were processed for western blot as before.[2,13,14] Briefly, protein samples (20 µg) were separated on a 10% SDS-PAGE and transferred to a PVDF membrane which was incubated overnight at 4°C with antibodies against Netrin-1 (1:1000, Abcam Inc, Toronto, ON, Canada) and a-Tubulin (1:20000, Cell Signaling, Danvers, MA, USA) for loading control. Protein bands were detected by chemiluminescence (Bio-Rad, Mississauga, ON, Canada) and analysed using Image Lab system software (Bio-Rad, Mississauga, ON, Canada).

Data Analysis

Planned comparisons were made between treatment groups for each experiment. Sex as a biological variable was included as a between-subjects factor when appropriate. Neuroanatomical data were analyzed using two-way mixed-design ANOVAs with genotype as a between-subjects factor and subregion as a within-subjects factor. To quantify the complexity of PFC neuron dendritic arbors, we used the Dendritic Complexity Index (DCI).[15] Behavioral data from the CPP test was analysed using Student's t-tests or one-way ANOVA. Data from the Go/No-Go task across sessions was analysed using Generalized Estimating Equations (GEE), while area under the curve was assessed using Student's t-tests or two-way ANOVA. Sigmoidal curve fitting was performed in MATLAB, by fitting commission errors to session with a sigmoid of the form $y = Min+(Max-Min)/ 1+10^{\wedge}(x50-x)*p$ where Min is the lower asymptote, Max is the upper asymptote, x50 is the position parameter denoting the training day at which the slope of the curve is maximal, and p determines the steepness of the sigmoid curve. The resulting fit was used to derive an index of improvement in the commission errors, defined as the day sustaining a half-maximal rate of commission errors (M50). For qPCR experiments, Student's t-tests or one-way ANOVA were used to assess treatment effect. When representations of data were normalized for graphs, all statistics were performed on the raw data. When post hoc testing was used, the most appropriate correction for multiple comparisons was chosen based on the factor design of the ANOVA in order to maximize power and not violate statistical assumptions. A Tukey's multiple comparisons test was used when all samples were independent and all possible interactions were considered. Sidak's multiple comparisons test was used when all samples were independent but comparisons were made only within one factor of interest, as comparing all groups was redundant or irrelevant. Dunnett's multiple comparisons test was used when conditions were compared to an explicit control condition, such as the *LacZ* sgRNA construct which should not amplify endogenous genes in the CRISPRa experiments. Detailed information about all statistical tests used are presented in Tables 1-5 for the main figures, and Supplementary Tables 2-4 for Extended data figures. All statistical analyses were carried out using Prism software (GraphPad), with the exception of the GEE, which was done in SPSS.

***Extended data Figures and Legends***

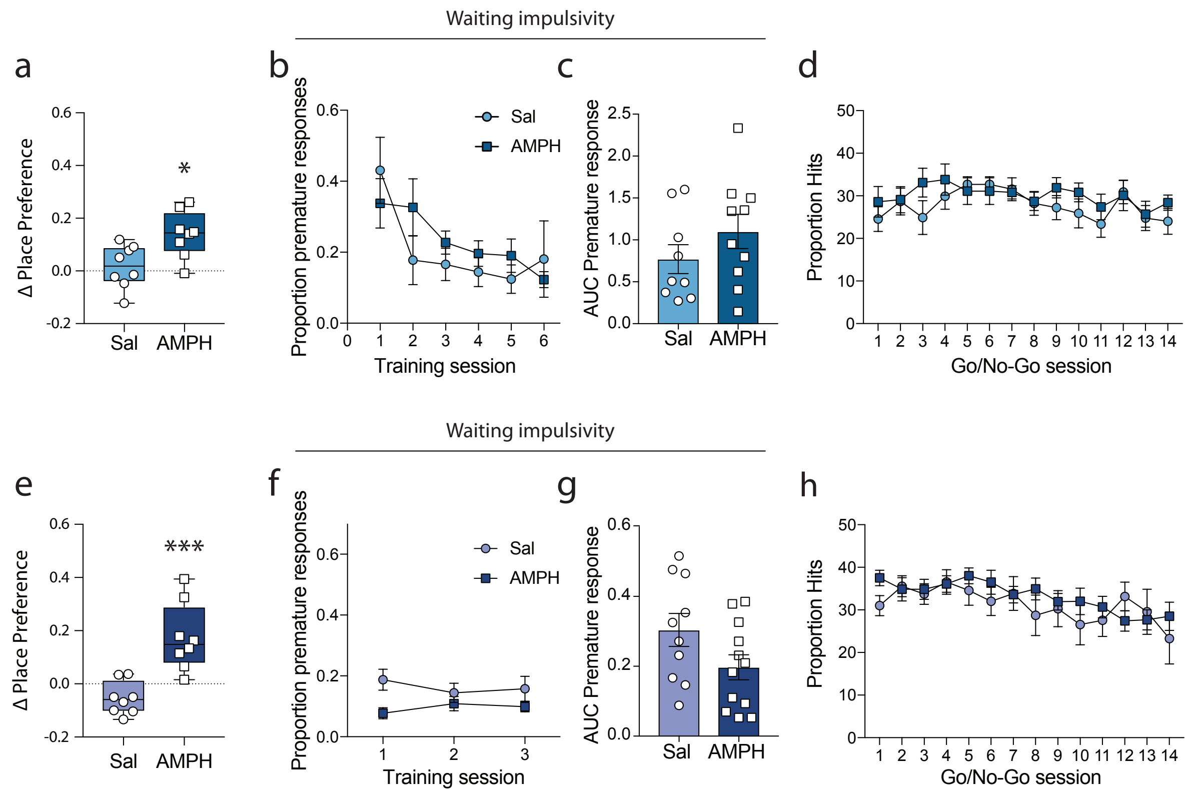

**Extended data figure 2** *Adolescent females are protected from the enduring effects of recreational-like amphetamine.* (a) Female mice show robust place preference to a recreational-like dose of 4 mg/kg amphetamine (AMPH) when tested in early adolescence (P21±1 to P31±1; Extended data Table 2A). (b-c) Adult female mice treated with AMPH in early adolescence do not show impairments in waiting impulsivity, measured by the level of premature responses during the final training phase of the Go/No-Go task over training days (b) or when assessed by the area under the curve (c) (Extended data Table 2B,C). (d) Adult female mice treated with AMPH in early adolescence do not show impairments in the correct response to 'Go' trials within Go/No-Go task (Extended data Table 2D). (e) Female mice show robust place preference to AMPH when tested in mid-adolescence (P35±1 to P44±1; Extended data Table 2E). (f-g) Adult female mice treated with AMPH in mid-adolescence do not show impairments in waiting impulsivity, measured by the level of premature responses during the final training phase of the Go/No-Go task over training days (f) or when assessed by the area under the curve (g) (Extended data Table 2F,G). (h) Adult female mice treated with AMPH in mid-adolescence do not show impairments in the correct response to 'Go' trials within Go/No-Go task (Extended data Table 2H).

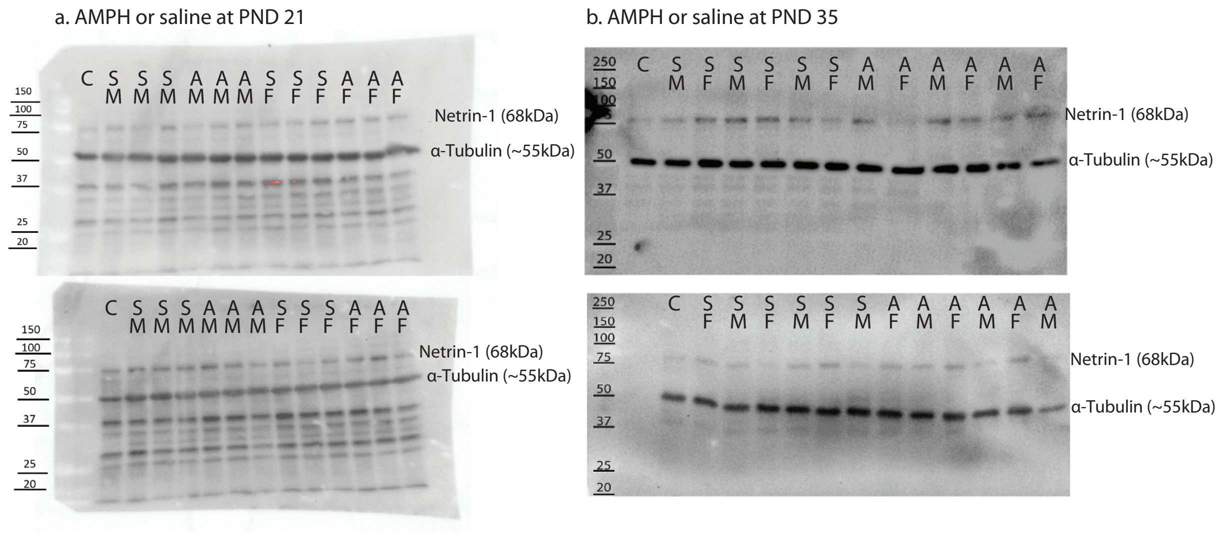

**Extended data figure 3** *Western blots for netrin-1 in the NAc.* Full blots for quantifications shown in Figure 3. (a) Western blot films from male (M) and female (F) mice treated with saline (S) or AMPH (A) in early adolescence (P21±1 to P31±1). (b) Western blot films from male (M) and female (F) mice treated with saline (S) or AMPH (A) in mid-adolescence (P35±1 to P44±1). Netrin-1 bands (68kDa) and α-tubulin reference bands (~55kDa) are indicated for each blot.

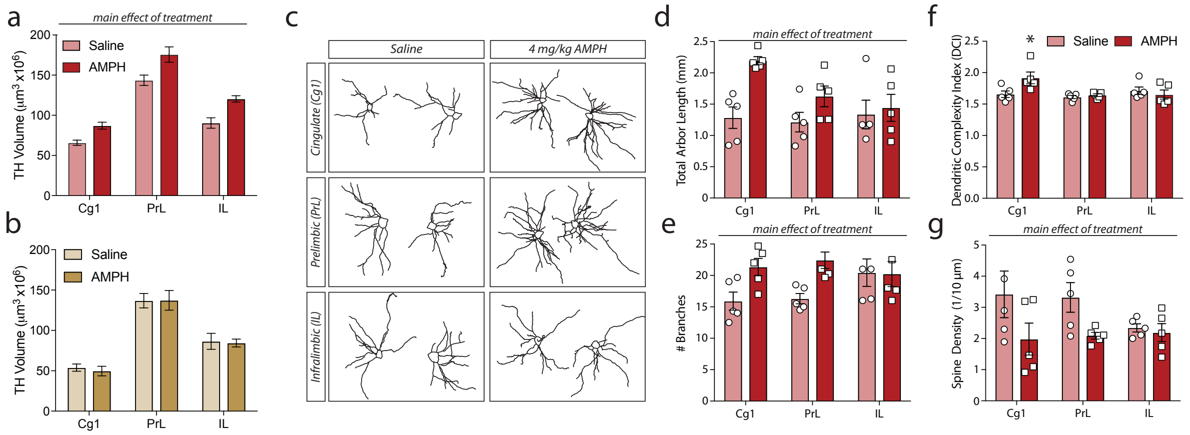

**Extended data figure 4** *Recreational-like AMPH increases dopamine innervation to the PFC and induces enduring changes in pyramidal neuron structure in male mice.* (a) Adult PFC dopamine input volume, measured by TH+ staining to the inner layers of three subregions of the medial PFC (Cg1, PrL, IL) is increased following exposure to recreational-like AMPH in early adolescent (P21±1 to P31±1) male mice (Extended data Table 3A). (b) Exposure to the therapeutic-like ALD in early adolescent (P21±1 to P31±1) mice does not produce enduring alterations in mPFC dopamine innervation volume (Extended data Table 3B). (c) Examples of Neurolucida tracings of PFC golgi-impregnated pyramidal neurons in adult male mice following treatment with saline or AMPH in early adolescence. (d-g) Neurolucida quantification of mPFC pyramidal neuron arbors in adult male mice treated with AMPH or saline in early adolescence. AMPH-treated mice show increases in total arbor length (d, Extended data Table 3C) and number of branches (e, Extended data Table 3D) across the three subregions of the mPFC studied. An increase in the complexity of pyramidal neuron arbors was evident only in the Cg1 subregion (f, Extended data Table 3E), while spine density assessed on tertiary branches was decreased across the PFC (g, Extended data Table 3F).

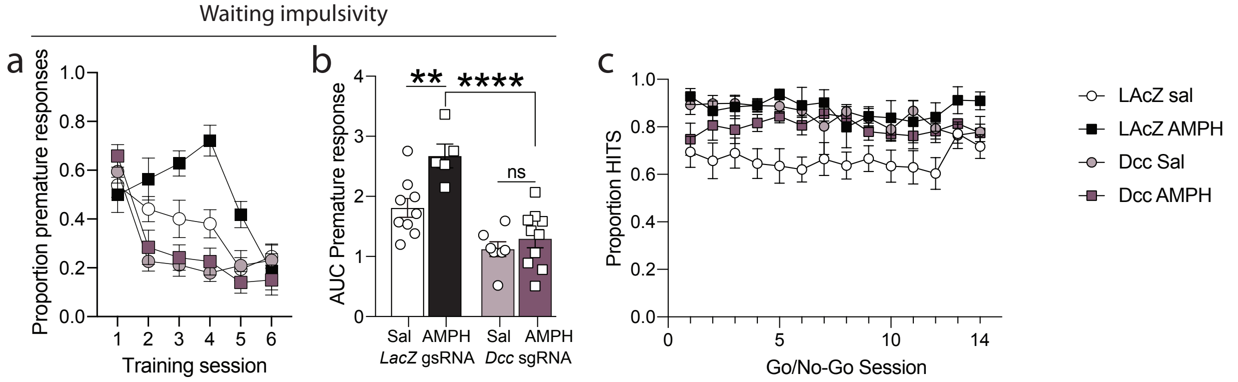

**Extended data figure 5** *CRISPRa upregulation of Dcc expression in the VTA prevents AMPH-induced deficits in waiting impulsivity.* (a-b). Adult male mice that received the *LacZ* sgRNA and were treated with AMPH in early adolescence showed impairments in waiting impulsivity, as measured by a high level of premature responses made over training days during the final training phase of the Go/No-Go task. In contrast, mice that received the *LacZ* sgRNA and were treated with saline and with mice that received the *Dcc* sgRNA and were treated with AMPH or with saline show significant improvement starting on the second training session (a, Extended data Table 4A). Assessment of the area under the curve (AUC, b, Extended data Table 4B), revealed that LacZ-injected mice show a significant difference in AUC between Saline and AMPH treated mice, but that this difference is abolished in the mice treated with the *Dcc* sgRNA. (d) No drug x construct x session effect is evident in the proportion of correct response to ‘Go’ trials (Hits) within the Go/No-Go task (Extended data Table 4C).

### Extended Data Table 1 : CRISPR RNA (crRNA) sequences for constructing single guide RNAs (sgRNAs)

| | Starting base pair | Target sequence | sgRNA sense | sgRNA anti-sense |
|---|---|---|---|---|
| **crRNA 1** | chr18:72351162 | GGCCTAGCGAAGCTTAGGGGGGG | CACCGGCCTAGCGAAGCTTAGGGG | AAACCCCCTAAGCTTCGCTAGGCC |
| **crRNA 2** | chr18:72351390 | GGTGTTTCACATAGGGCAAGTGG | CACCGGTGTTTCACATAGGGCAAG | AAACCTTGCCCTATGTGAAACACC |
| **crRNA 3** | chr18:72351532 | CTCGCGTTTGTTTTCCGTGTGGG | CACCCTCGCGTTTGTTTTCCGTG | AAACCACGGAAAACAAACGCGAGC |
| **crRNA 4** | chr18:72351809 | GTTAACACACTCTCATCATACGG | CACCGTTAACACACTCTCATCATA | AAACTATGATGAGAGTGTGTTAAC |

### Extended Data Table 2: Detailed statistics for Extended Data Figure 2

| | Statistical test | Factor | n | Statistic | 95% confidence interval | p value (adjusted where appropriate) | corresponding figure |
|---|---|---|---|---|---|---|---|
| **A** | | | **8/group** | | | | **Extended data Figure 2a** |
| | unpaired t test | Saline vs AMPH CPP | | t(14) =2.868 | 0.03082 to 0.2136 | **0.0124** | |
| **B** | | | **Saline = 5 AMPH = 6** | | | | **Extended data Figure 2b** |
| | Two-way mixed ANOVA | Interaction | | F (5, 45) = 0.7547 | | 0.5871 | |
| | | Training session (within subject) | | F (5, 45) = 4.586 | | **0.0018** | |
| | | Treatment (between subjects) | | F (1, 9) = 0.3616 | | 0.5625 | |
| **C** | | | **Saline = 5 AMPH = 6** | | | | **Extended data Figure 2c** |
| | unpaired t test | AUC | | t(17) = 1.219 | -0.2409 to 0.8999 | 0.2396 | |
| **D** | | | **Saline = 9 AMPH = 10** | | | | **Extended data Figure 2d** |
| | Two-way mixed ANOVA | Interaction | | F (13, 221) = 0.7133 | | 0.7493 | |

| | | | | | | | |
|---|---|---|---|---|---|---|---|
| | | Go/No-Go session (within subject) | | F (13, 221) = 1.782 | | **0.0471** | |
| | | Treatment (between subjects) | | F (1, 17) = 0.9632 | | 0.3401 | |
| **E** | | | **8/group** | | | | **Extended data Figure 2e** |
| | | Saline vs AMPH CPP | | t(14) =4.55 | 0.1206 to 0.3357 | **0.0005** | |
| **F** | | | **Saline = 8 AMPH = 12** | | | | **Extended data Figure 2f** |
| | Two-way mixed ANOVA | Interaction | | F (2, 36) = 2.067 | | 0.1413 | |
| | | Training session (within subject) | | F (2, 36) = 0.04828 | | 0.9529 | |
| | | Treatment (between subjects) | | F (1, 18) = 4.76 | | **0.0426** | |
| **G** | | | **Saline = 8 AMPH = 12** | | | | **Extended data Figure 2g** |
| | unpaired t test | AUC | | t(20)=1.842 | -0.2282 to 0.01416 | 0.0803 | |
| **H** | | | **Saline = 7 AMPH = 12** | | | | **Extended data Figure 2h** |
| | Two-way mixed ANOVA | Interaction | | F (13, 221) = 1.408 | | 0.1569 | |
| | | Go/No-Go session (within subject) | | F (13, 221) = 4.575 | | **<0.0001** | |

| | | | | | |
|---|---|---|---|---|---|
| | | Treatment (between subjects) | | $F_{(1, 17)} = 0.3712$ | 0.5504 |

### Extended Data Table 3: Detailed statistics for Extended Data Figure 4

| | **Statistical test** | **Factor** | **n** | **Statistic** | **95% confidence interval** | **p value (adjusted where appropriate)** | **corresponding figure** |
|---|---|---|---|---|---|---|---|
| **A** | | | **Saline = 5 AMPH = 4** | | | | **Extended data Figure 4a** |
| | Two-way mixed ANOVA | Interaction | | F (2, 14) = 1.002 | | 0.3921 | |
| | | Subregion (within subject) | | F (2, 14) = 221.9 | | **<0.0001** | |
| | | Treatment (between subjects) | | F (1, 7) = 15.07 | | **0.006** | |
| **B** | | | **Saline = 4 AMPH = 6** | | | | **Extended data Figure 4b** |
| | Two-way mixed ANOVA | Interaction | | F (2, 16) = 0.07807 | | 0.9252 | |
| | | Subregion (within subject) | | F (2, 16) = 111.2 | | **<0.0001** | |
| | | Treatment (between subjects) | | F (1, 8) = 0.03607 | | 0.8541 | |
| **C** | | | **5/group** | | | | **Extended data Figure 4d** |
| | Two-way mixed ANOVA | Interaction | | F (2, 16) = 3.008 | | 0.0778 | |
| | | Subregion (within subject) | | F (2, 16) = 2.754 | | 0.0938 | |
| | | Treatment (between subjects) | | F (1, 8) = 9.272 | | **0.0159** | |

| | | | | | | | |
|---|---|---|---|---|---|---|---|
| **D** | | | **5/ group** | | | | **Extended data Figure 4e** |
| | Two-way mixed ANOVA | Interaction | | F (2, 16) = 3.255 | | 0.0651 | |
| | | Subregion (within subject) | | F (2, 16) = 0.8069 | | 0.4636 | |
| | | Treatment (between subjects) | | F (1, 8) = 5.544 | | **0.0463** | |
| **E** | | | **5/ group** | | | | **Extended data Figure 4f** |
| | Two-way mixed ANOVA | Interaction | | F (2, 16) = 4.064 | | **0.0374** | |
| | | Subregion (within subject) | | F (2, 16) = 3.904 | | **0.0416** | |
| | | Treatment (between subjects) | | F (1, 8) = 2.245 | | 0.1724 | |
| | Sidak's multiple comparisons test | Saline vs. AMPH within Cg1 | | t(24) = 3.09 | 0.04403 to 0.475 | **0.0149** | |
| | Sidak's multiple comparisons test | Saline vs. AMPH within PrL | | t(24) = 0.3517 | -0.186 to 0.245 | 0.9799 | |
| | Sidak's multiple comparisons test | Saline vs. AMPH within IL | | t(24) = 0.7362 | -0.2773 to 0.1537 | 0.85 | |
| **F** | | | **5/ group** | | | | **Extended data Figure 4g** |
| | Two-way mixed ANOVA | Interaction | | F (2, 16) = 1.226 | | 0.3195 | |
| | | Subregion (within subject) | | F (2, 16) = 0.6534 | | 0.5336 | |
| | | Treatment (between subjects) | | F (1, 8) = 6.85 | | **0.0308** | |

### Extended Data Table 4: Detailed statistics for Extended Data Figure 5

| | Statistical test | Factor | n | Statistic | 95% confidence interval | p value (adjusted where appropriate) | corresponding figure |
|---|---|---|---|---|---|---|---|
| A | | | ***LacZ* + Saline= 9 LacZ + AMPH= 5 Dcc + Saline = 8 *Dcc* + AMPH = 10** | | | | **Extended data Figure 5a** |
| | Generalized Estimating Equations analysis (GEE) | Drug (between subjects) | | Wald Chi-Square = 5.408 (df = 1) | | **0.02** | |
| | | Construct (between subjects) | | Wald Chi-Square = 25.669 (df = 1) | | **<0.001** | |
| | | Session (within subjects) | | Wald Chi-Square = 127.826 (df = 5) | | **<0.001** | |
| | | Drug x Construct | | Wald Chi-Square = 4.484 (df = 1) | | **0.034** | |
| | | Construct x Session | | Wald Chi-Square = 84.410 (df = 5) | | **<0.001** | |
| | | Drug x Session | | Wald Chi-Square = 27.922 (df = 5) | | **<0.001** | |
| | | Drug x Construct x Session | | Wald Chi-Square = 22.721 (df = 5) | | **<0.001** | |
| B | | | ***LacZ* + Saline= 9 LacZ + AMPH= 5 Dcc + Saline = 8** | | | | **Extended data Figure 5b** |

| | | | | | | | |
|---|---|---|---|---|---|---|---|
| | | | ***Dcc* + AMPH = 10** | | | | |
| | Two-way ANOVA | Interaction | | F (1, 27) = 4.39 | | **0.0457** | |
| | | sgRNA construct (between subject) | | F (1, 27) = 39.45 | | ***<0.0001*** | |
| | | Drug (between subject) | | F (1, 27) = 9.991 | | **0.0039** | |
| | Tukey's multiple comparisons test | LacZ:Sal vs. LacZ:AMPH | | q(27) = 4.959 | -1.538 to -0.1897 | **0.0082** | |
| | Tukey's multiple comparisons test | **LacZ:Sal vs. Dcc:Sal** | | q(27) = 4.372 | **0.0789 to 1.297** | **0.0224** | |
| | Tukey's multiple comparisons test | LacZ:Sal vs. Dcc:AMPH | | q(27) = 3.574 | -0.04244 to 1.068 | *0.0781* | |
| | Tukey's multiple comparisons test | LacZ:AMPH vs. Dcc:Sal | | q(27) = 8.487 | 0.8439 to 2.259 | **<0.0001** | |
| | Tukey's multiple comparisons test | LacZ:AMPH vs. Dcc:AMPH | | q(27) = 8.049 | 0.7145 to 2.038 | **<0.0001** | |
| | Tukey's multiple comparisons test | Dcc:Sal vs. Dcc:AMPH | | q(27) = 1.138 | -0.7705 to 0.4203 | 0.8516 | |
| **C** | | | ***LacZ* + Saline= 9**<br>**LacZ + AMPH= 5**<br>**Dcc + Saline = 8**<br>***Dcc* + AMPH = 10** | | | | **Extended data Figure 5c** |
| | Generalized Estimating Equations analysis (GEE) | Drug (between subjects) | | Wald Chi-Square = 5.74 (df = 1) | | **0.17** | |
| | | Construct (between subjects) | | Wald Chi-Square = 2.292 (df = 1) | | **0.13** | |
| | | Session (within subjects) | | Wald Chi-Square = 15.95 (df = 13) | | **0.252** | |

| | | | | | |
|---|---|---|---|---|---|
| | | Drug x Construct | | Wald Chi-Square = 13.703 (df = 1) | **<0.001** |
| | | Construct x Session | | Wald Chi-Square = 21.309 (df = 13) | **0.67** |
| | | Drug x Session | | Wald Chi-Square = 62.54 (df = 13) | **<0.001** |
| | | Drug x Construct x Session | | Wald Chi-Square = 17.60 (df = 13) | **0.173** |

**Supplementary References**

1. Cuesta, S. *et al.* DCC-related developmental effects of abused- versus therapeutic-like amphetamine doses in adolescence. *Addict Biol* e12791 (2019) doi:10.1111/adb.12791.

2. Reynolds, L. M. *et al.* DCC Receptors Drive Prefrontal Cortex Maturation by Determining Dopamine Axon Targeting in Adolescence. *Biol Psychiat* 83, 181 192 (2018).

3. Reynolds, L. M. *et al.* Quantifying Dopaminergic Innervation in Rodents Using Unbiased Stereology. in *Dopaminergic System Function and Dysfunction: Experimental Approaches* (eds. Evans, J. A. F. & Vargas, P. F. H.) vol. 193 31–63 (Springer, 2022).

4. Paxinos, G. & Franklin, K. B. J. *The Mouse Brain in Stereotaxic Coordinates*. (Academic Press, 2008).

5. Savell, K. E. *et al.* A Neuron-Optimized CRISPR/dCas9 Activation System for Robust and Specific Gene Regulation. *Eneuro* 6, ENEURO.0495-18.2019 (2019).

6. Labun, K., Montague, T. G., Gagnon, J. A., Thyme, S. B. & Valen, E. CHOPCHOP v2: a web tool for the next generation of CRISPR genome engineering. *Nucleic Acids Res* 44, W272–W276 (2016).

7. Montague, T. G., Cruz, J. M., Gagnon, J. A., Church, G. M. & Valen, E. CHOPCHOP: a CRISPR/Cas9 and TALEN web tool for genome editing. *Nucleic Acids Res* 42, W401–W407 (2014).

8. Platt, R. J. *et al.* CRISPR-Cas9 Knockin Mice for Genome Editing and Cancer Modeling. *Cell* 159, 440–455 (2014).

9. Fasano, C., Thibault, D. & Trudeau, L. Culture of Postnatal Mesencephalic Dopamine Neurons on an Astrocyte Monolayer. *Curr Protoc Neurosci* 44, 3.21.1-3.21.19 (2008).

10. Manitt, C. *et al.* The netrin receptor DCC is required in the pubertal organization of mesocortical dopamine circuitry. *J Neurosci* 31, 8381 8394 (2011).

11. Gibb, R. & Kolb, B. A method for vibratome sectioning of Golgi–Cox stained whole rat brain. *J Neurosci Meth* 79, 1–4 (1998).

12. Cuesta, S. *et al.* Non-Contingent Exposure to Amphetamine in Adolescence Recruits miR-218 to Regulate Dcc Expression in the VTA. *Neuropsychopharmacol* 43, 900 911 (2018).

13. Grant, A. *et al.* Netrin-1 receptor-deficient mice show enhanced mesocortical dopamine transmission and blunted behavioural responses to amphetamine. *Eur J Neurosci* 26, 3215 3228 (2007).

14. Cuesta, S. *et al.* Dopamine Axon Targeting in the Nucleus Accumbens in Adolescence Requires Netrin-1. *Frontiers Cell Dev Biology* 8, 487 (2020).

15. Lom, B. & Cohen-Cory, S. Brain-Derived Neurotrophic Factor Differentially Regulates Retinal Ganglion Cell Dendritic and Axonal Arborization In Vivo. *J Neurosci* 19, 9928–9938 (1999).
